# Xylose phosphatase activity of dystroglycan self-regulates its receptor function

**DOI:** 10.1101/2025.10.10.681626

**Authors:** Ishita Chandel, David Venzke, Bailey A. Wollesen, Liping Yu, Kevin P. Campbell

## Abstract

Dystroglycan (DG), a transmembrane receptor crucial for tissue development and pathogen entry, harbors a glycan composed of xylose and glucuronic acid called matriglycan. Loss of matriglycan or reduction in its length affects DG function, causing dystroglycanopathies. However, the mechanism underlying matriglycan extension is unknown. Here, we show that a Golgi xylose kinase facilitates initiation of matriglycan synthesis by transiently adding a phosphate to the xylose of matriglycan primer. Matriglycan extends when the phosphate is removed from xylose by the N-terminal domain of dystroglycan (DGN). DGN has the DXDXT/V motif found in haloacid dehalogenase (HAD) domains of hydrolases, conditional mutations in which reduce matriglycan length and cause disease in mice. Our work reveals an unexpected glycan phosphatase function of DG in regulating matriglycan extension on itself.

## Main Text

Dystroglycan (DG) is a ubiquitously expressed cell membrane receptor required for the development of brain (*1*), muscles (*2, 3*) and heart (*4, 5*) in multicellular organisms. The extracellular part of DG, called alpha DG (α-DG) is extensively glycosylated on its mucin-like domain (317-485) (*6*). Glycosylated α-DG is hijacked by Old World arenaviruses such as Lassa fever virus, lymphocytic choriomeningitis virus (LCMV) (*7*) and the pathogenic bacterium *Mycobacterium leprae* (*8*), for entry into host cells. The most crucial and complex glycosylation on α-DG is the phosphorylated O-mannose trisaccharide [GalNAcβ1,3-GlcNAcβ1,4-(phosphate-6) Man-O-Ser] called core M3 glycan (*6*) (Fig. 1A). The core M3 phosphorylated trisaccharide is elongated twice with ribitol-5-phosphate (Rbo5P) (*9, 10*) after which the enzymes Ribitol xylosyltransferase 1 (RXYLT1) (*11, 12*) and beta-1,4-glucuronyltransferase1 (B4GAT1) (*13*) sequentially add a disaccharide unit [GlcA-β1,4-Xyl], creating the LARGE1 primer (Fig. 1A). This primed structure is recognized by the bifunctional glycosyltransferase like-acetylglucosaminyltransferase 1 (LARGE1), that further adds alternating xylose and glucuronic acid units (*14*) [-GlcA-β1,3-Xyl-α1,3]_n_ to the core M3 glycan (Fig. 1A). The linear repeating structure of [-GlcA-β1,3-Xyl-α1,3]_n_ is termed matriglycan (*15*).

**Fig. 1.**
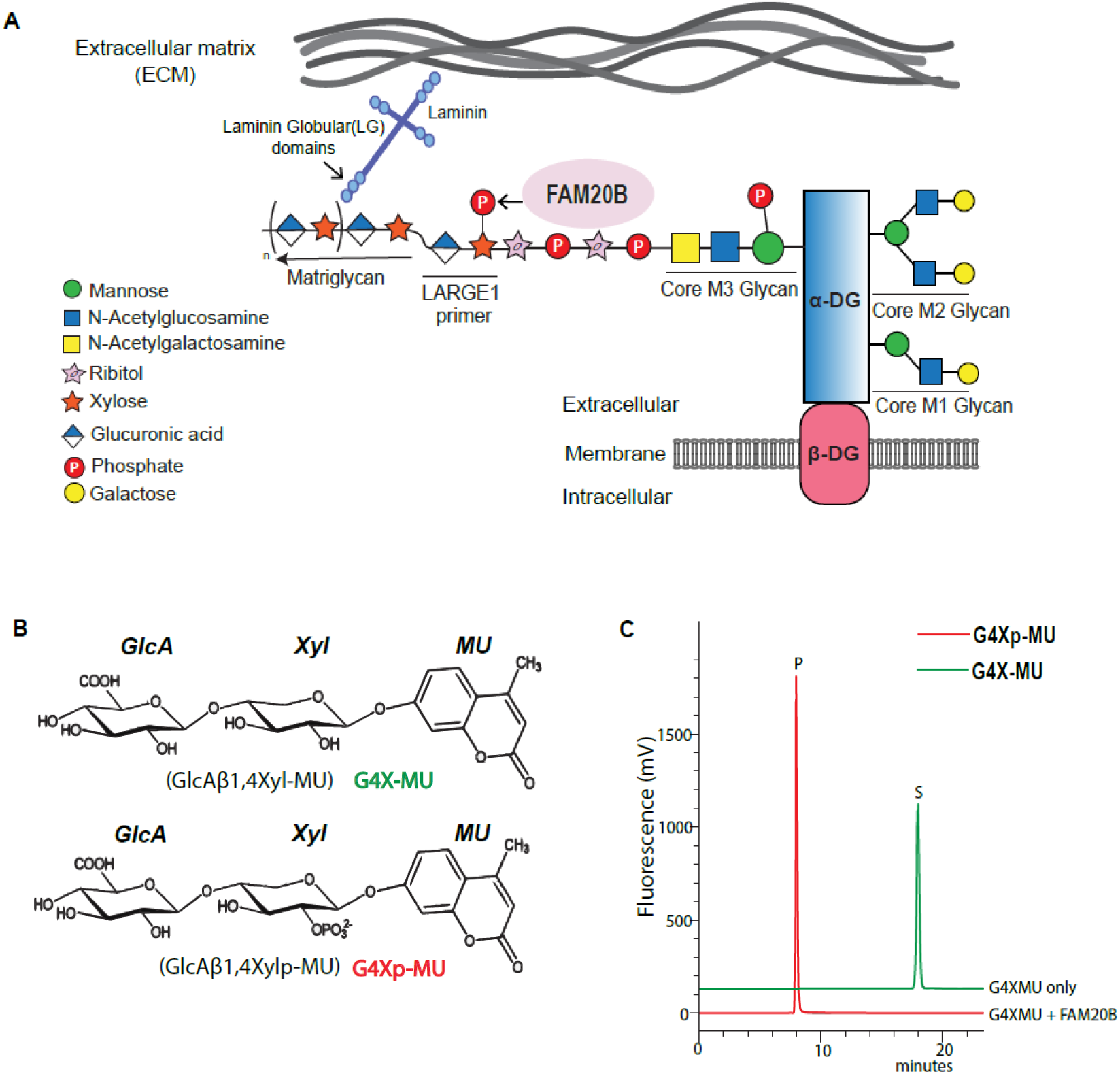
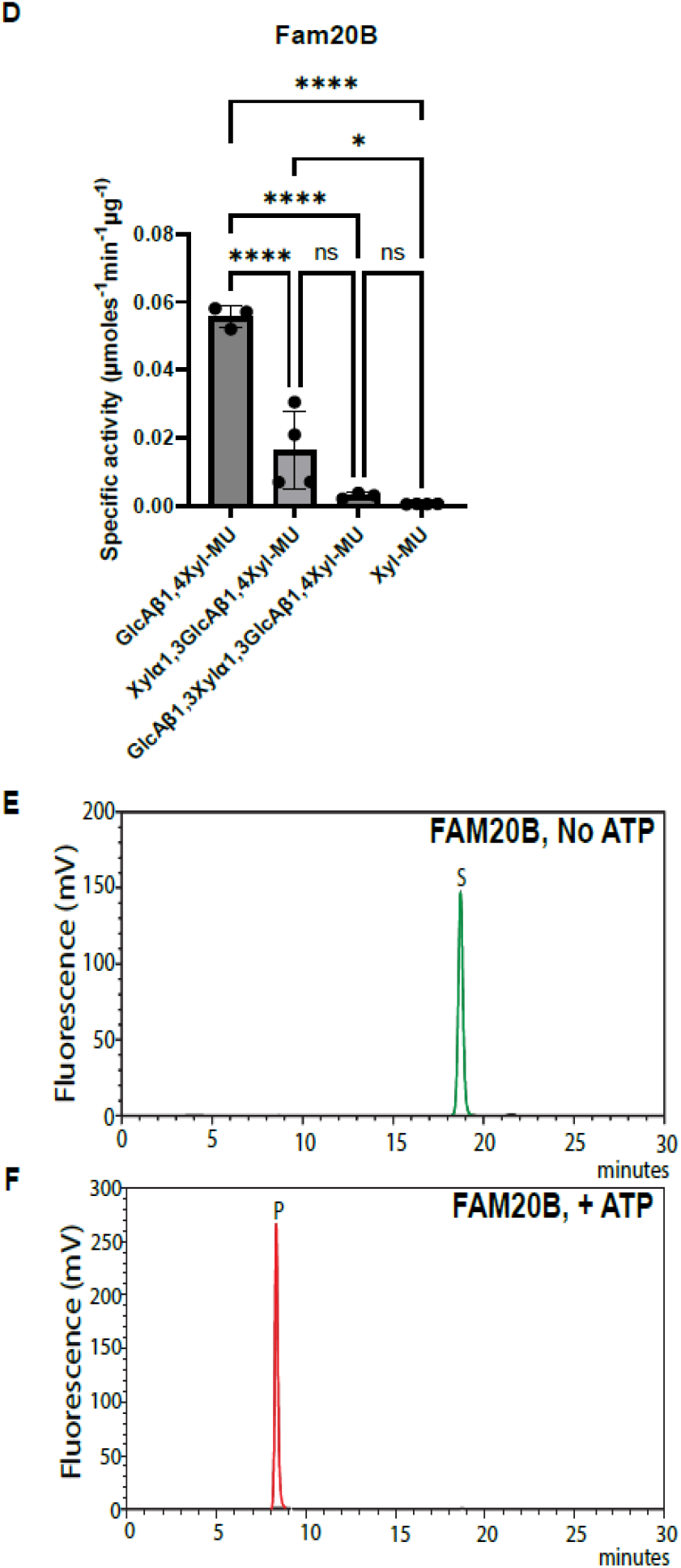
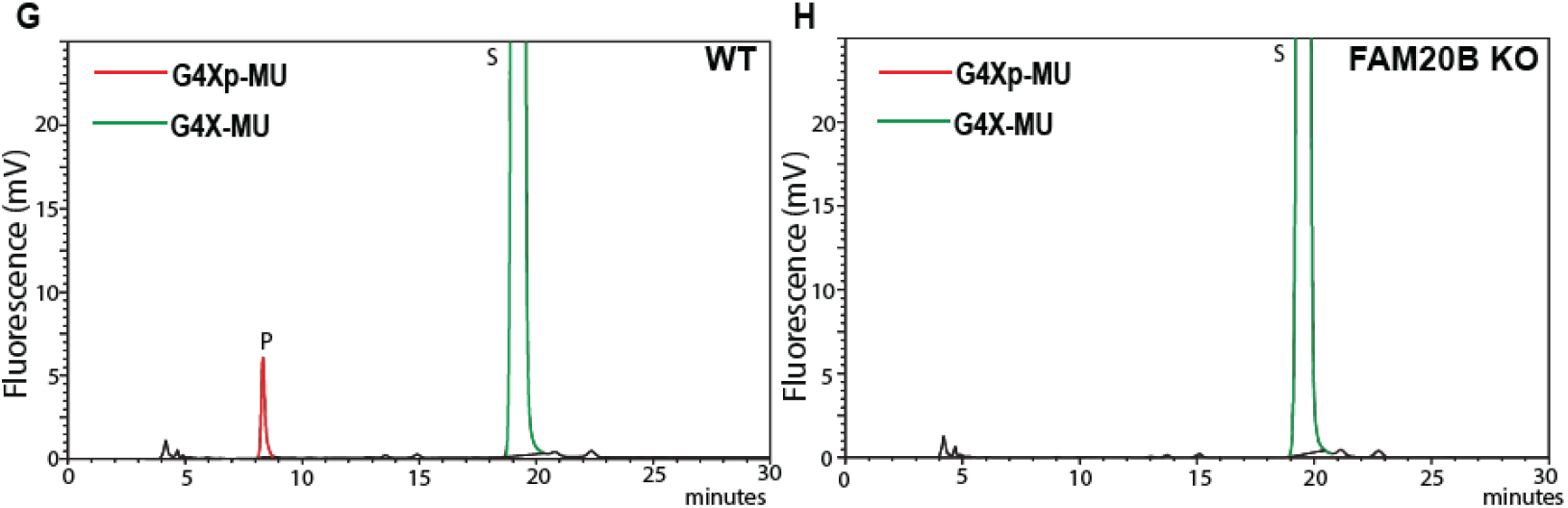
FAM20B phosphorylates xylose of matriglycan primer disaccharide. (**A**) Cartoon depicts FAM20B adding the phosphate to xylose in the LARGE1 primer disaccharide as part of the core M3 mannose modification on dystroglycan (α-DG). (**B**) Structure of chemically synthesized primer disaccharide on methylumbelliferone (MU), G4X-MU and its phosphorylated form, G4Xp-MU. (**C**) Substrate (S) G4X-MU is converted to product (P) G4Xp-MU in presence of FAM20B and separated on a C-18 reverse phase column. (**D**) Specific activity of FAM20B with different sugars is calculated from independent reactions. Statistical significance was determined by Ordinary one-way ANOVA was used with Tukey’s post-hoc test (*p value < 0.05, ****p value < 0.0001, ns= not significant). (**E, F**) FAM20B mediated conversion of substrate (S) G4X-MU to product (P) G4Xp-MU (**E**) without ATP and (**F**) with ATP. (**G, H**) Chromatogram of substrate (S), G4XMU to product (P), G4XpMU conversion in the presence of (**G**) WT HAP cell lysate and (**H**) FAM20B KO HAP cell lysate.

Pathogens such as Lassa fever virus, LCMV and *M. leprae* bind to matriglycan to enter host cells (*16-18*). Matriglycan enables DG to bind to laminin globular (LG) domains of extracellular matrix (ECM) proteins to maintain DG receptor function and thus structural integrity of cells (*19*). Matriglycan alone can recapitulate DG binding and function (*20*). Mutations in any of the 18 genes (*21*) required to produce the matriglycan containing core M3 structure leads to muscular dystrophies with or without brain abnormalities and cardiomyopathy, such as muscle-eye-brain disease (MEB) and Walker-Warburg syndrome (*22*). Severity of disease is dependent on the length of matriglycan (*19*), and even though a short chain of matriglycan can bind to LG domains, it cannot prevent disease (*21, 23*). The length of matriglycan is regulated during development and is cell-type specific (*19*). However, the mechanism by which matriglycan length is regulated is unknown. α-DG is composed of three distinct domains: the N-terminal (α-DGN) domain (29-316), a central mucin-like domain (317-485) and a C-terminal domain (486-653). A patient mutation in α-DGN (T192M) was reported to reduce matriglycan length and cause muscular dystrophy and cognitive impairment (*24*). Removal of α-DGN in mouse muscles significantly reduces the length of matriglycan, causing severe muscle pathology (*23*). Control of matriglycan length is suggested to be intrinsic to α-DGN (*25*). α-DGN is also thought to serve as an intracellular substrate recognition site for LARGE1 (*26*). However, this has never been shown and the mechanism by which α-DGN controls matriglycan length remains unclear.

Here, we show that α-DGN acts as a xylose phosphatase to regulate matriglycan extension. LARGE1 synthesizes matriglycan using the primer disaccharide unit [GlcA-β1,4-Xyl] as substrate. We show that a Golgi glycosaminoglycan xylosylkinase, Family with sequence similarity 20, member B (FAM20B), an atypical secretory kinase, adds a phosphate to the xylose of the primer disaccharide. LARGE1 then binds to the phosphorylated primer with higher affinity to initiate matriglycan synthesis. LARGE1 makes matriglycan at a faster rate when using the phosphorylated primer disaccharide, indicating it is a preferred substrate for initiating matriglycan synthesis. However, the phosphate on xylose must be removed by α-DGN for matriglycan extension to occur. Muscle-specific mutations in the potential active site of α-DGN lead to a shorter matriglycan and consequently, muscular dystrophy in mice. Thus, our results demonstrate that matriglycan extension is a fine-tuned process regulated by primer xylose phosphorylation and dephosphorylation, which is critical for DG to perform its physiological functions and prevent disease.

### FAM20B phosphorylates xylose of matriglycan primer

FAM20B is a glycosaminoglycan xylosylkinase that facilitates extension of glycosaminoglycan (GAG) chains by adding phosphate to xylose of tetrasaccharide linkage (*27, 28*). Deletion of FAM20B causes embryonic lethality in mice (*29*). To test if FAM20B can add a phosphate to the xylose of matriglycan primer disaccharide (recognized by LARGE1), we chemically synthesized the disaccharide on a xylose-β-methylumbelliferone (X-MU) using recombinant B4GAT1, hence creating the primed structure [GlcA-β1,4-Xyl-β-MU] (*13*) recognized by LARGE1 (Fig. 1B). From here on, we will refer to the primer disaccharide as G4X-MU and its phosphorylated form as G4Xp-MU.

Next, we incubated G4X-MU with recombinant FAM20B and separated the products using C18 column chromatography. FAM20B was able to phosphorylate xylose in an ATP-dependent manner and showed the highest activity for the disaccharide as substrate (Fig 1C-F). FAM20B did not phosphorylate the xylose of LARGE1-synthesized matriglycan oligosaccharides: Xylα1,3-GlcAβ1,3-Xyl-α1,3-GlcA-β-MU; GlcAβ1,3-Xylα1,3-GlcAβ1,3-Xylα1,3-GlcA-β-MU (G4-MU and G5-MU respectively) (*30*) (fig.S1A to S1D). FAM20B belongs to a family of kinases (*31*), of which FAM20C was also shown to phosphorylate xylose of the proteoglycan tetrasaccharide *in vitro* (*32*). Therefore, we tested several Golgi-associated kinases (FAM20C, FAM198A, and Fjx-1) for xylose phosphorylation activity, none of which showed activity towards the xylose in the primer disaccharide (G4X-MU) (fig. S1E to S1G). To further confirm if native endogenous FAM20B can phosphorylate xylose of G4X-MU, we performed the phosphorylation assay using WT and FAM20B knockout (KO) cell lysates. The WT cell lysate showed phosphorylation of xylose, whereas the FAM20B KO did not (Fig. 1G and H NMR, we also tested the binding affinity of LARGE1 with the ). We were unable to detect FAM20B protein when we probed the cell lysate with anti-human FAM20B antibody, confirming that the KO cells lacked expression of this protein (fig. S1H and S1I).

FAM20B adds phosphate to the hydroxyl group of the second carbon of xylose in the proteoglycan tetrasaccharide linkage (*28*). To test if xylose of the primer disaccharide unit (G4X-MU) was also phosphorylated at the second position, we performed nuclear magnetic resonance (NMR) experiments on the phosphorylated product (G4Xp-MU) (Fig. 2A) purified by analytical C-18 column chromatography. The ^1^H and ^13^C resonances of the product were assigned by using heteronuclear multiple quantum coherence (HMQC) and heteronuclear 2-bond correlation (H2BC) spectra (Fig. 2B). The 1D ^31^P NMR spectrum showed one phosphate signal originating from G4Xp-MU (Fig. 2C) which gave a strong COSY peak to β-Xyl H2 (Fig. 2D) indicating that the phosphate was linked to the 2-position of the xylose. These results indicate that FAM20B is a matriglycan primer xylosylkinase.

**Fig. 2.**
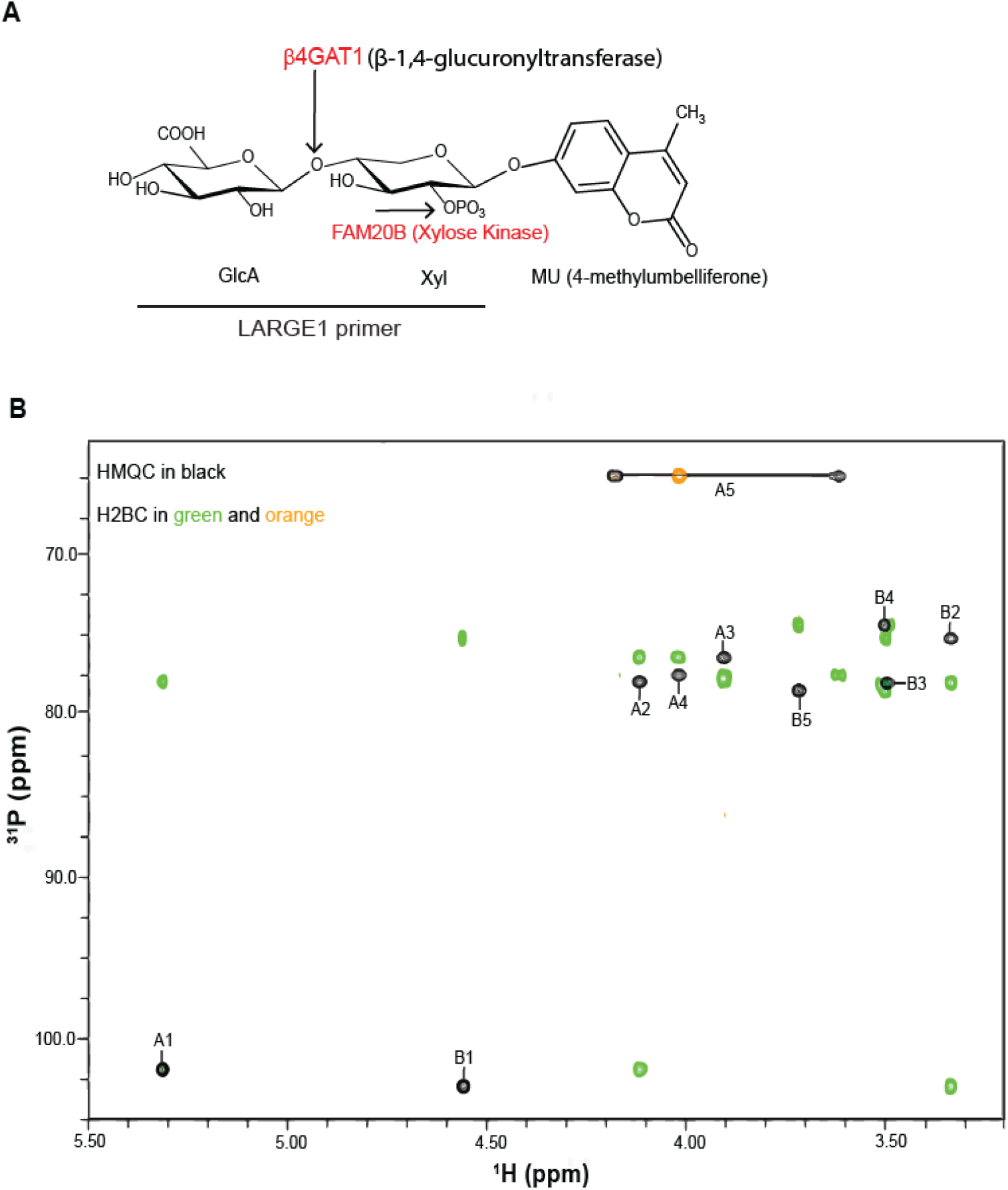
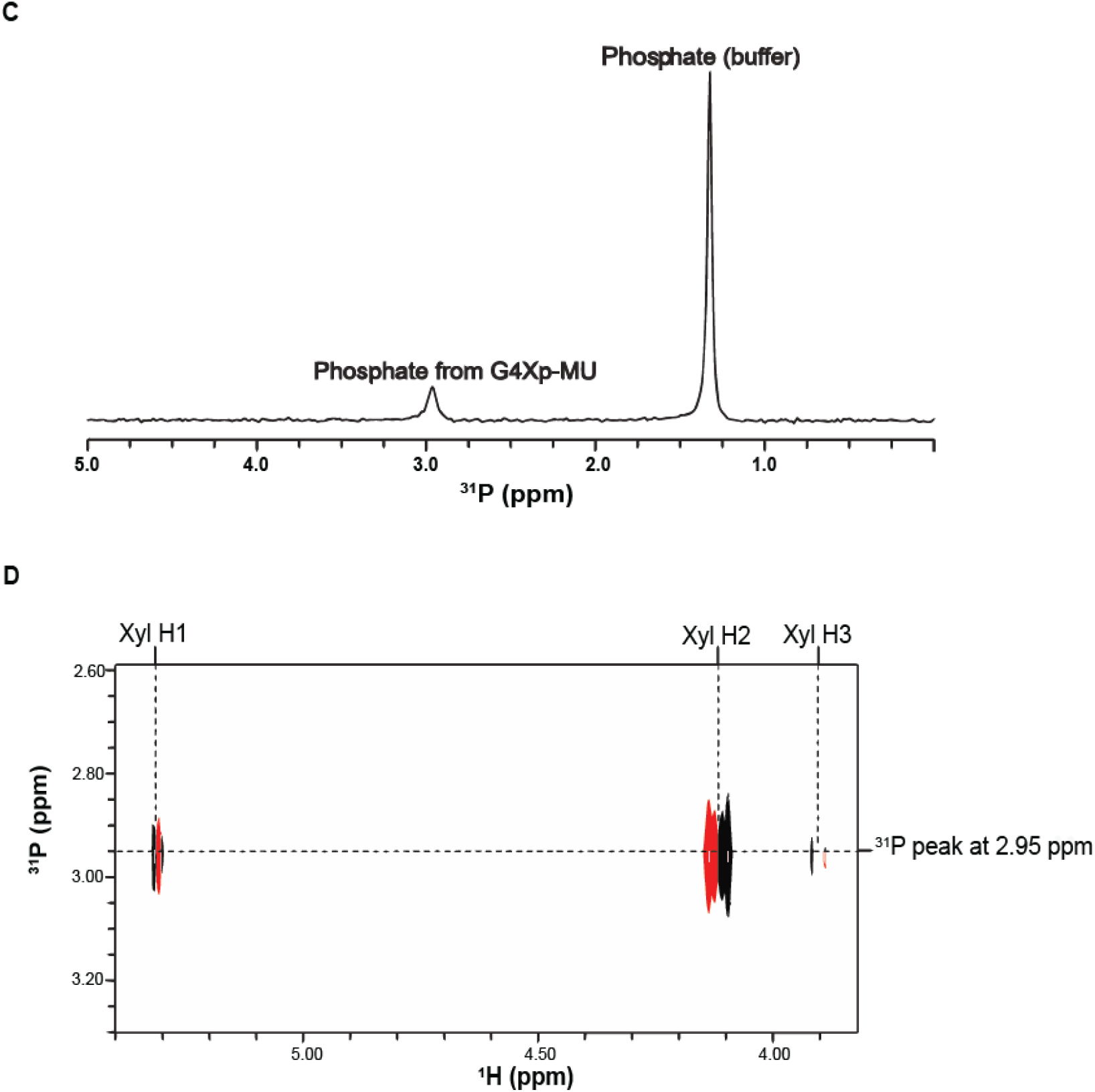
FAM20B phosphorylates hydroxyl group at the second carbon of xylose. (**A**) Cartoon depicting FAM20B phosphorylates at position 2 of β-Xyl of G4X-MU. B4GAT1: β-1,4-glucuronic acid transferase 1. FAM20B: atypical secretory kinase family with sequence similarity 20, member B. (**B**) NMR assignments of ^13^C/^1^H HMQC and H2BC spectra of G4Xp-MU are shown. The HMQC cross peaks are labeled with a capital letter followed by a number. The letter A or B represents residue A for Xyl or B for GlcA, whereas the number indicates the position on that sugar residue. (**C**) 1D ^31^P NMR spectrum, showing one phosphate signal from G4Xp-MU at 2.95 ppm. (**D**) The ^31^P peak at 2.95 ppm observed in the 1D ^31^P NMR spectrum gives a strong COSY peak to β-Xyl H2 and two very weak COSY peaks to β-Xyl H1 and H3, indicating that the phosphate group is attached to β-Xyl at position 2.

### FAM20B-mediated xylose phosphorylation facilitates matriglycan initiation

To test whether phosphorylation of primer xylose affects matriglycan synthesis by LARGE1, we performed a LARGE1 enzyme assay (*14*) using the phosphorylated and unphosphorylated primer disaccharide (G4Xp-MU and G4X-MU, respectively) as substrate. LARGE1 produced more matriglycan and thus showed higher specific activity when using G4Xp-MU substrate as compared to G4X-MU (Fig. 3A-G). Using NMR, we also tested the binding affinity of LARGE1 with the two substrates. LARGE1 exhibited higher binding affinity towards G4Xp-MU than G4X-MU (Fig. 3H-J). These data suggest that xylose phosphorylation promotes matriglycan initiation.

**Fig. 3.**
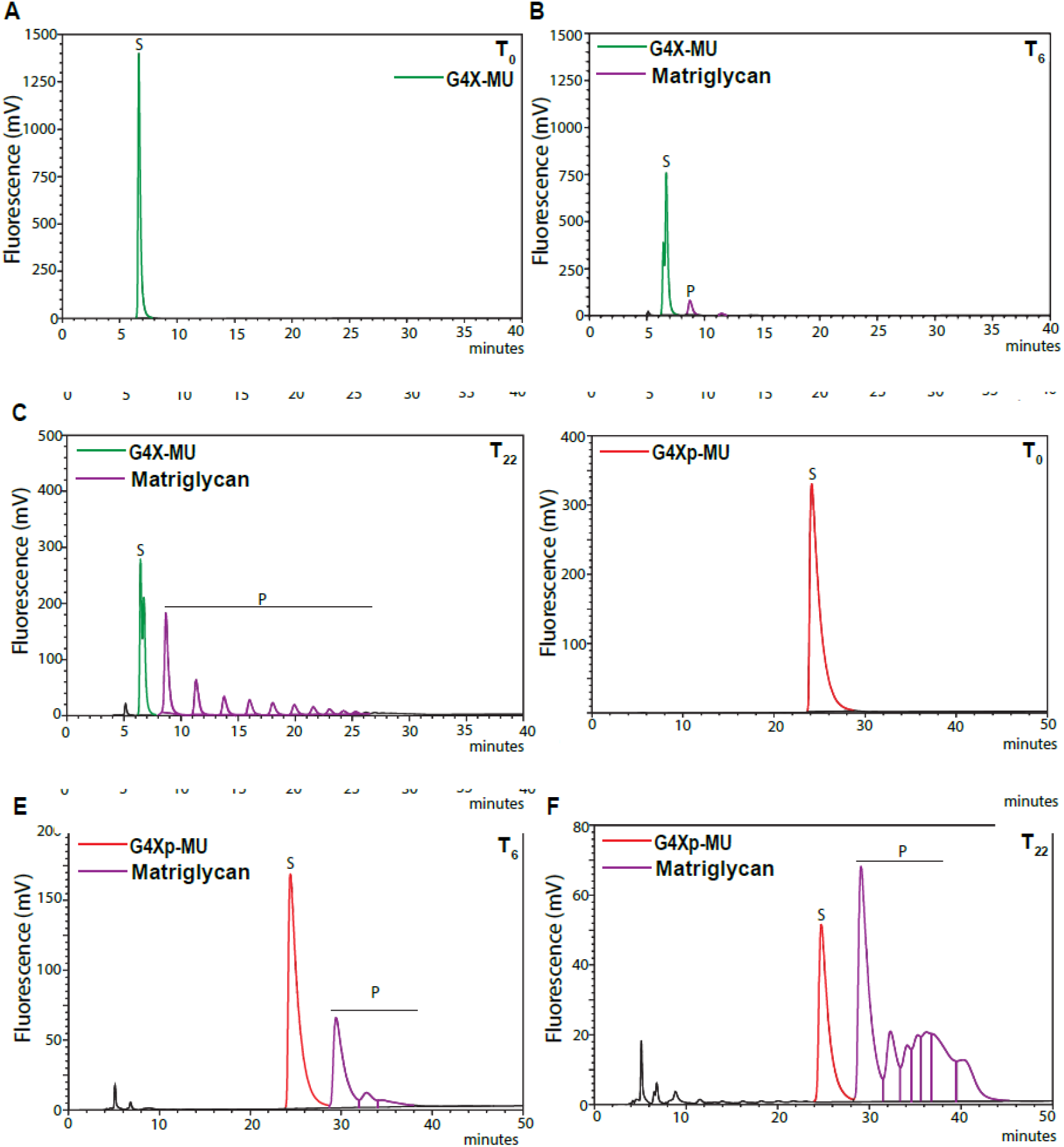
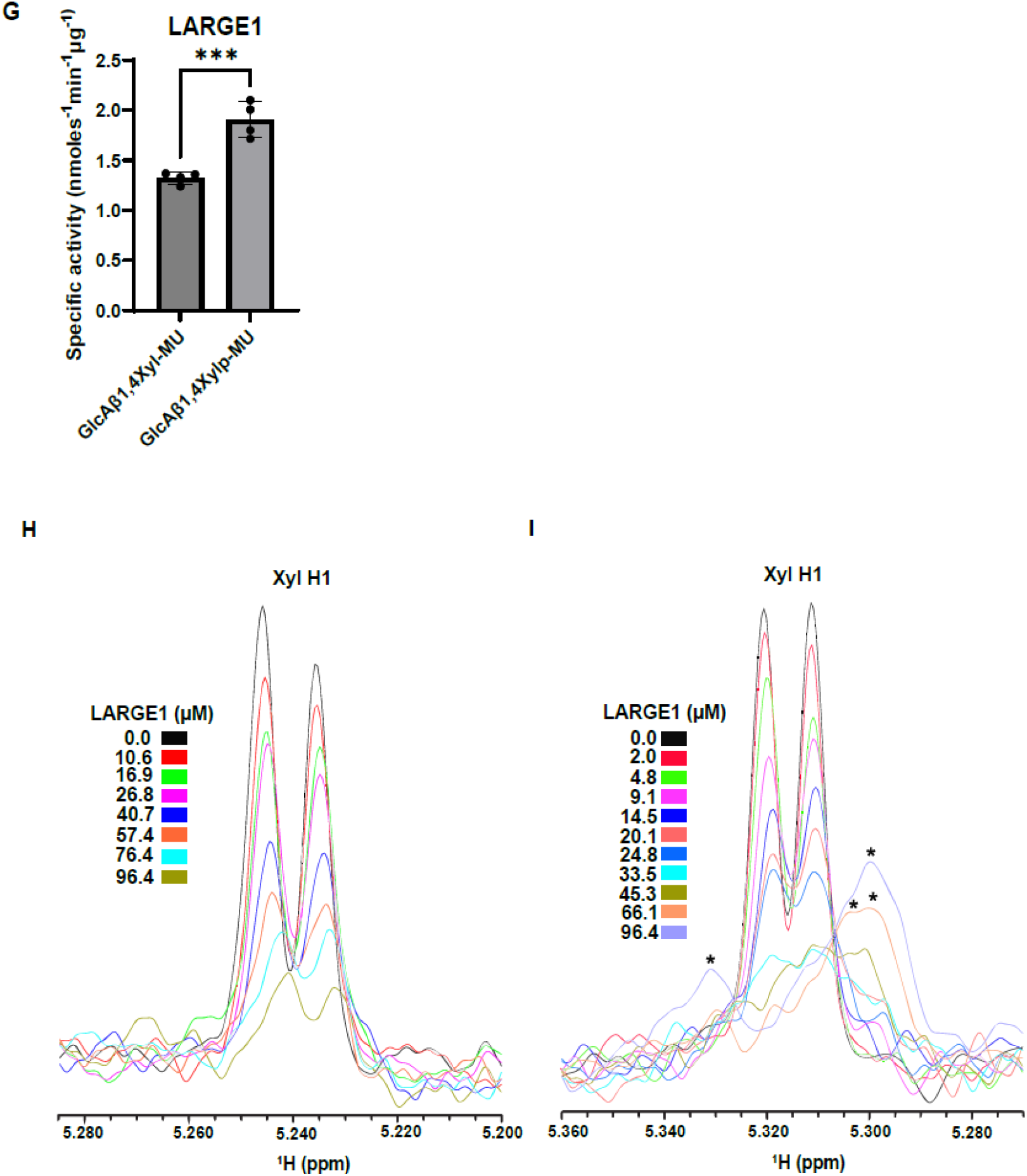
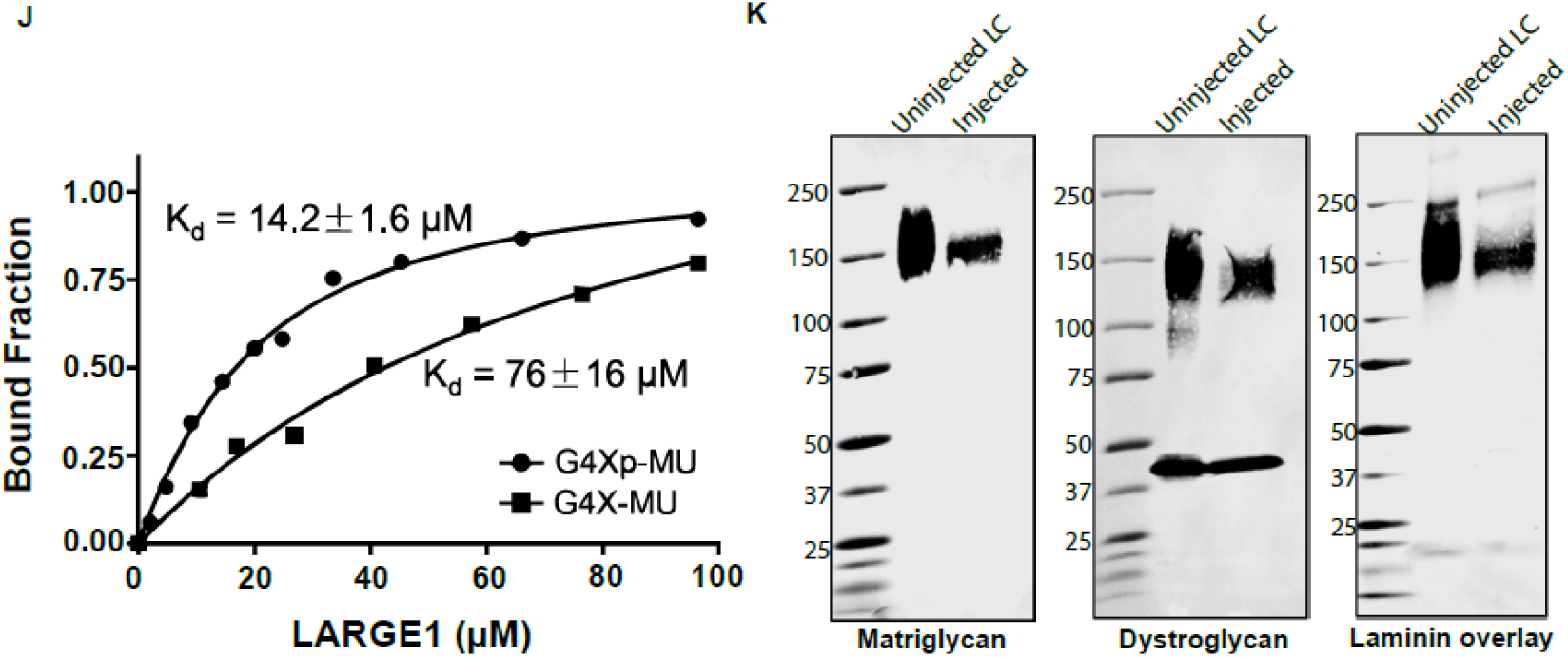
FAM20B xylose phosphorylation facilitates matriglycan initiation. Timed reaction was done to produce matriglycan with G4X-MU as a substrate at (**A**) 0 hours, (**B**) 6 hours, (**C**) 22 hours, and with G4Xp-MU as a substrate at (**D**) 0 hours, (**E**) 6 hours, (**F**) 22 hours. Products are separated on a SAXS anion exchange column. Multiple homogenous peaks depict matriglycan. (**G**) LARGE1 specific activity calculated using G4X-MU and G4Xp-MU is shown. Statistical significance determined by Student’s unpaired two-tailed t-test (***p-value<0.001). (**H, I**) 1D ^1^H NMR spectra of the anomeric region of (**H**) G4X-MU and (**I**) G4Xp-MU were acquired for the glycan concentration of 10.0 µM in the presence of various concentrations of LARGE1 as indicated. The peak Xyl H1 is derived from the residue Xyl anomeric H1 proton. Black stars mark the peaks derived from the protein, especially at higher protein concentrations as confirmed by using the apo protein sample without glycan. (**J**) Fitting of the NMR binding data by calculating the bound fraction through measuring the difference in the peak intensity of the Xyl H1 anomeric proton in the absence (free form) and presence (bound form) of mammalian LARGE1, then divided by the peak intensity of the free form. The data were then fitted using GraphPad Prism to obtain the dissociation constant K_d_. (**K**) Immunoblot analysis of heart from FAM20B knockdown mice. Glycoproteins were enriched using wheat-germ agglutinin (WGA)-agarose with 10 mM EDTA. Immunoblotting was performed to detect matriglycan (IIH6), core α-DG, β-DG (AF6868) (the broad 100-200 kDa band represents α-DG and the discrete ∼ 42kDa band represents β-DG), and laminin overlay. Molecular weights are indicated on the left of each blot.

To test if FAM20B phosphorylation is required to efficiently produce matriglycan *in vivo*, we created FAM20B knockdown mice since FAM20B deletion is embryonic lethal. We injected newborn mice with Ad5-U6-FAM20B shRNA using the retro-orbital (RO) sinus route for systemic delivery. FAM20B knockdown mice developed slowly and were smaller than their un-injected littermates (fig. S2A to S2E), exhibited hindlimb clasping and died around 3 weeks post-injection. Detailed investigations of muscle physiology in FAM20B knockdown mice could not be conducted due to early lethality. This is consistent with a previous report that showed FAM20B deletion caused severe stunted embryonic growth, delayed development and death at around E13.5 (*29*).

A previous study reported that adenovirus injections via the RO route lead to the majority of the adenovirus delivered to the heart (*33*). Therefore, we probed the heart of FAM20B knockdown mice with anti-FAM20B, matriglycan, and dystroglycan antibodies. Heart tissue of FAM20B knockdown mice showed a significant reduction in FAM20B protein levels (fig. S3A), a reduction in molecular weight of matriglycan and dystroglycan and reduced laminin binding (Fig. 3K). Skeletal muscles of FAM20B knockdown mice were also assessed however, they did not show a reduction in FAM20B protein levels (fig. S3A) or any change in the molecular weights of matriglycan or dystroglycan (fig. S3B), indicating that indeed the adenovirus was efficiently delivered to the heart but not muscles. Together, our *in vitro* and *in vivo* results provide strong evidence that FAM20B-mediated phosphorylation is required for efficient initiation of matriglycan.

### N-terminal domain of dystroglycan (α-DGN) exhibits xylose phosphatase activity

FAM20B-mediated xylose phosphorylation of the proteoglycan tetrasaccharide linkage is most likely transient (*34*) and is specifically required for the initiating/priming step in heparan sulfate biosynthesis (*35*). Therefore, we posited that FAM20B-mediated xylose phosphorylation of matriglycan primer is also transient, and α-DGN may be involved in dephosphorylation. To test this hypothesis, we conducted structural and sequence analysis of α-DGN, which revealed that it has a DXDXT/V motif (Fig. 4A and B) commonly found in the haloacid dehalogenase (HAD) domain of enzymes, which are a large superfamily of phosphohydrolases found in all three super kingdoms of life (*36*). To test whether α-DGN has phosphate hydrolysis activity, we expressed and purified recombinant α-DGN from a mammalian system (*37*). α-DGN was purified using a sequential two-step purification method: first using His-tag, followed by size exclusion chromatography (fig. S4A to S4E).

**Fig. 4.**
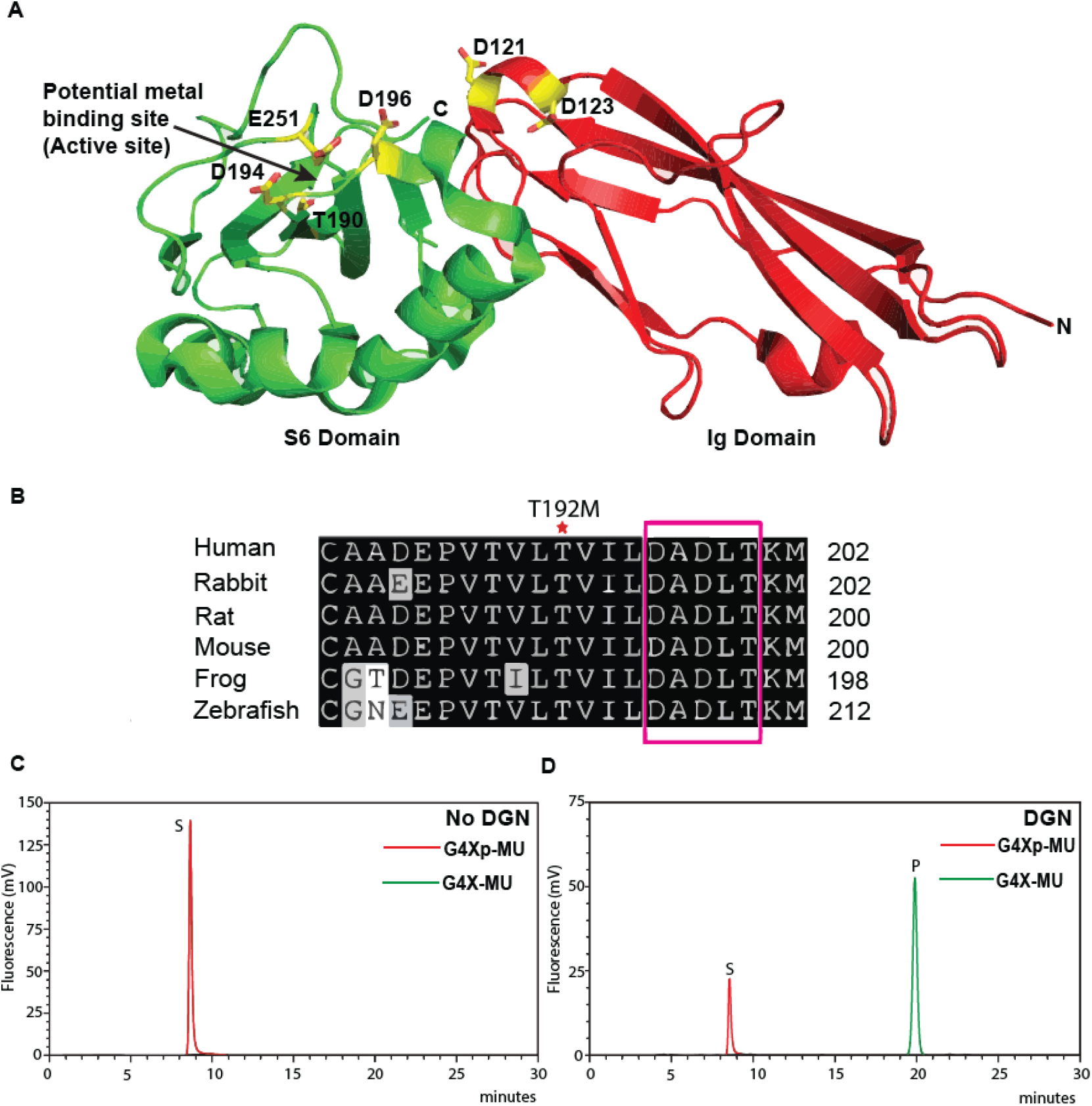
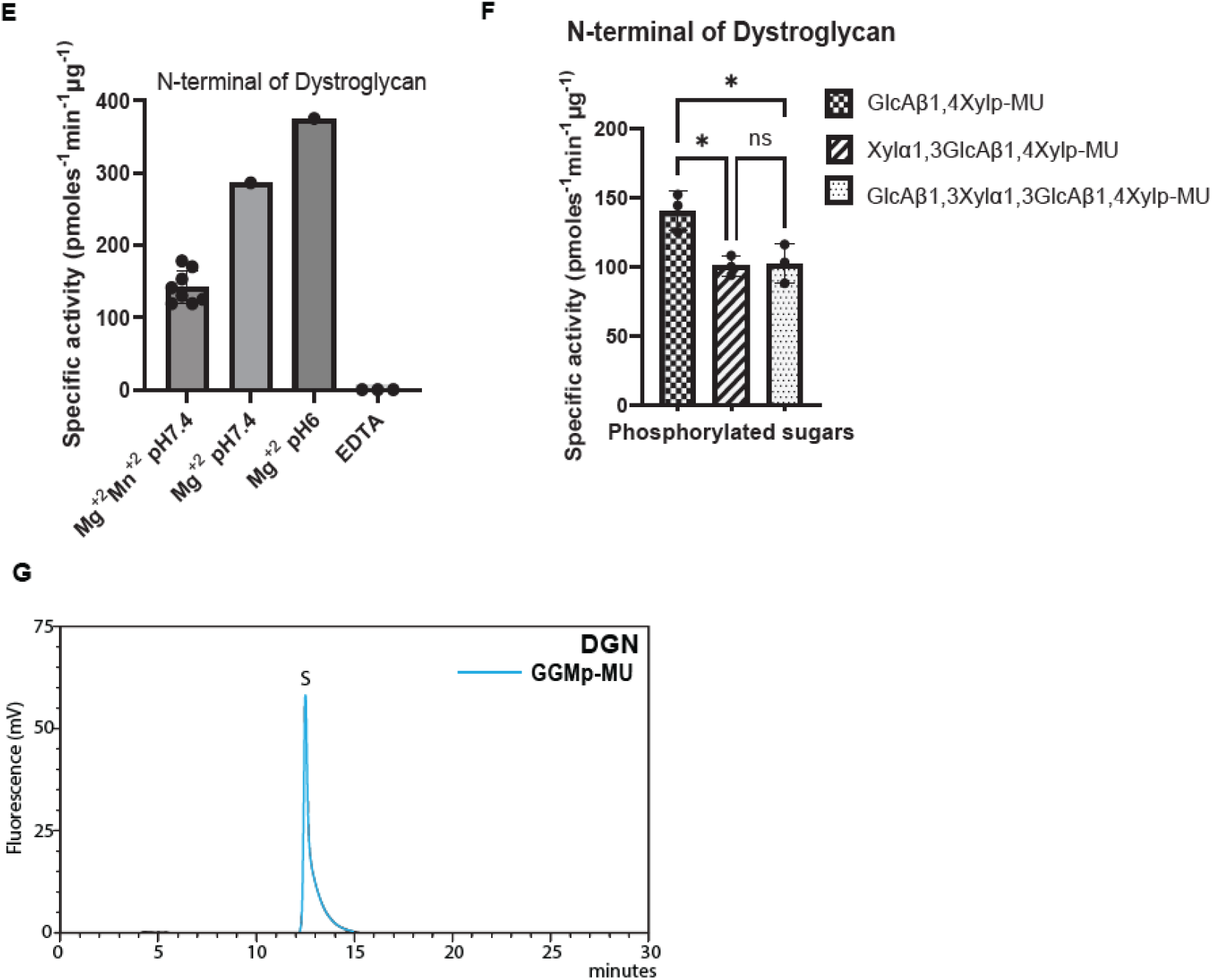
N-terminal domain of dystroglycan is a xylose phosphatase. (**A**) The structure of mouse α-dystroglycan N-terminal (α-DGN) contains two domains (Ig and S6). Amino acids D194, D196 and E251 of S6 domain are a potential metal-binding active site. Amino acid residues 161-178 are missing from the structure. (**B**) Conserved HAD domain motif in α-DGN (DXDXT/V) is highlighted in a pink box. Patient mutation T192M close to the HAD motif is marked with a red asterisk (*). (**C-D**) Chromatogram showing conversion of substrate (S) G4Xp-MU to product (P) G4X-MU in the presence of α-DGN. (**E**) Specific activity of α-DGN is calculated under different conditions from independent reactions. (**F**) Specific activity of α-DGN for glycans of different lengths is calculated from independent reactions. Statistical significance was determined by Ordinary one-way ANOVA used with Tukey’s post-hoc test (*p value < 0.05, ns= not significant). (**G**) Chromatogram showing α-DGN is unable to hydrolyze phosphate from phosphorylated mannose trisaccharide (GGMp-MU).

We incubated purified α-DGN with the phosphorylated primer disaccharide G4Xp-MU and separated the products using C18 column chromatography. α-DGN hydrolyzed the phosphate from G4Xp-MU in a metal-ion and pH dependent manner (Fig. 4C-E). α-DGN exhibited the highest activity towards the disaccharide, although it was moderately active towards the elongated tri- and tetrasaccharides (Fig. 4F).

Since DG has another phosphate modification on the mannose of the core M3 trisaccharide (Fig. 1A), which is added by the enzyme protein-O-mannose kinase (POMK) (*21, 38*), we wanted to test if α-DGN hydrolyzes that phosphate. To test this, we synthesized the phosphorylated mannose trisaccharide on a GlcNAc-β1,4-mannose-6-phosphate 4-MU using recombinant β-1,3-N-Acetylgalactosaminyltransferase (B3GALNT2), creating GalNAcβ1,3-GlcNAcβ1,4-(phosphate-6) Man-O-MU, from here on referred to as GGMp-MU. α-DGN failed to hydrolyze the phosphate on the mannose of GGMp-MU (Fig. 4G). This indicates that α-DGN exhibits phosphatase activity specifically towards the xylose phosphate of the LARGE1 primer disaccharide in the core M3 modification. Together, our results indicate that α-DGN acts as a HAD-like xylose phosphatase.

### Xylose phosphatase activity of α-DGN is required for matriglycan extension

HAD-like phosphatases have a characteristic N-terminal motif I, which contains two Asp residues (DXD) (*39*). We found two such motifs in α-DGN, one in the Ig domain at the D121/D123 residues and another in the S6 domain at D194/D196 residues of the mouse sequence (fig. S5A). To test whether these motifs are required for matriglycan extension, we mutated the Asp residues to Asn using CRISPR (D121N/D123N and D194N/D196N). Mice with homozygous D194N/D196N mutations were embryonic lethal (Fig. 5A), whereas the mice with D121N/D123N mutations survived and did not show changes in the molecular weight of DG and matriglycan or physiological changes (fig. S5B to S5E).

**Fig. 5.**
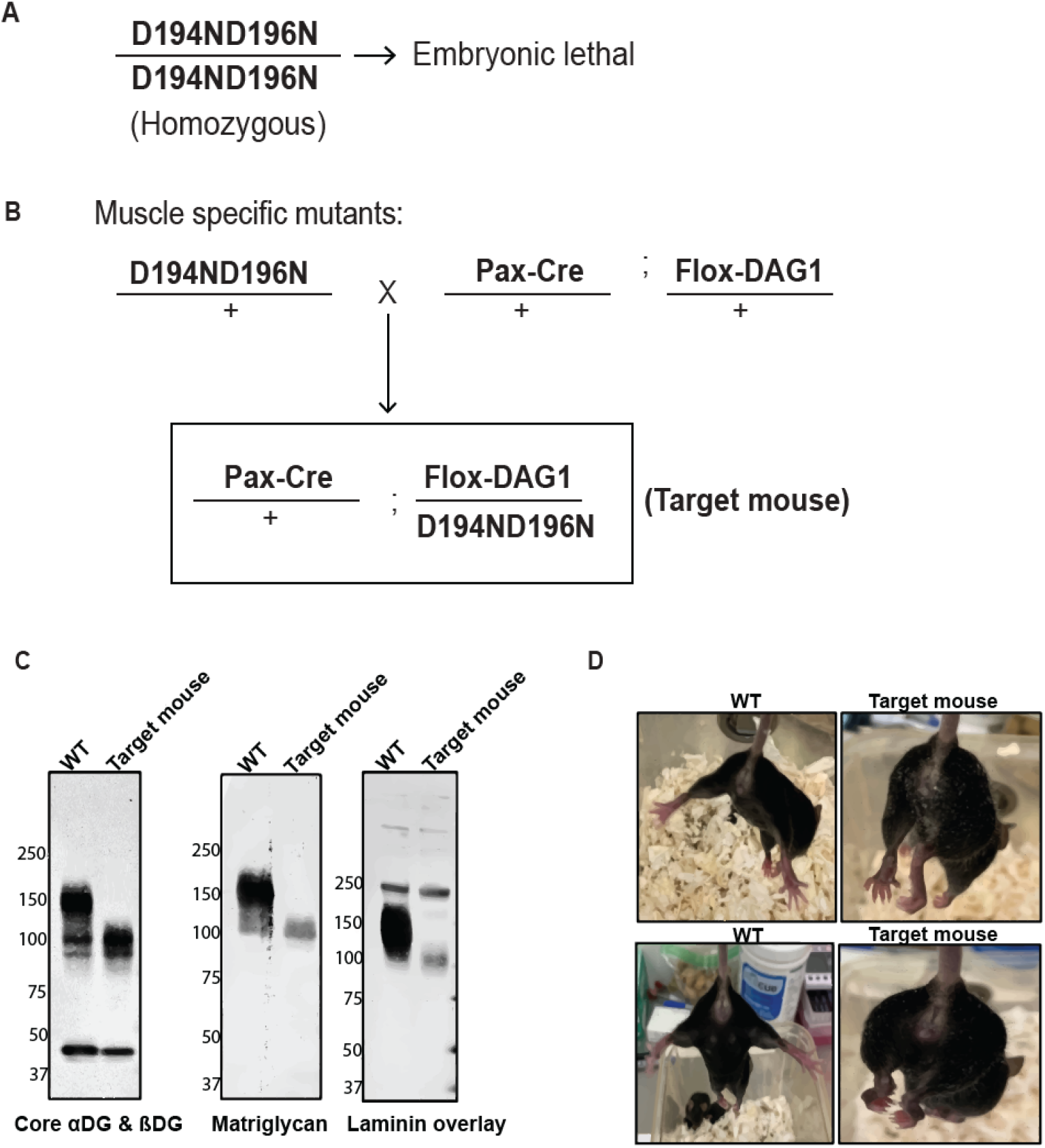
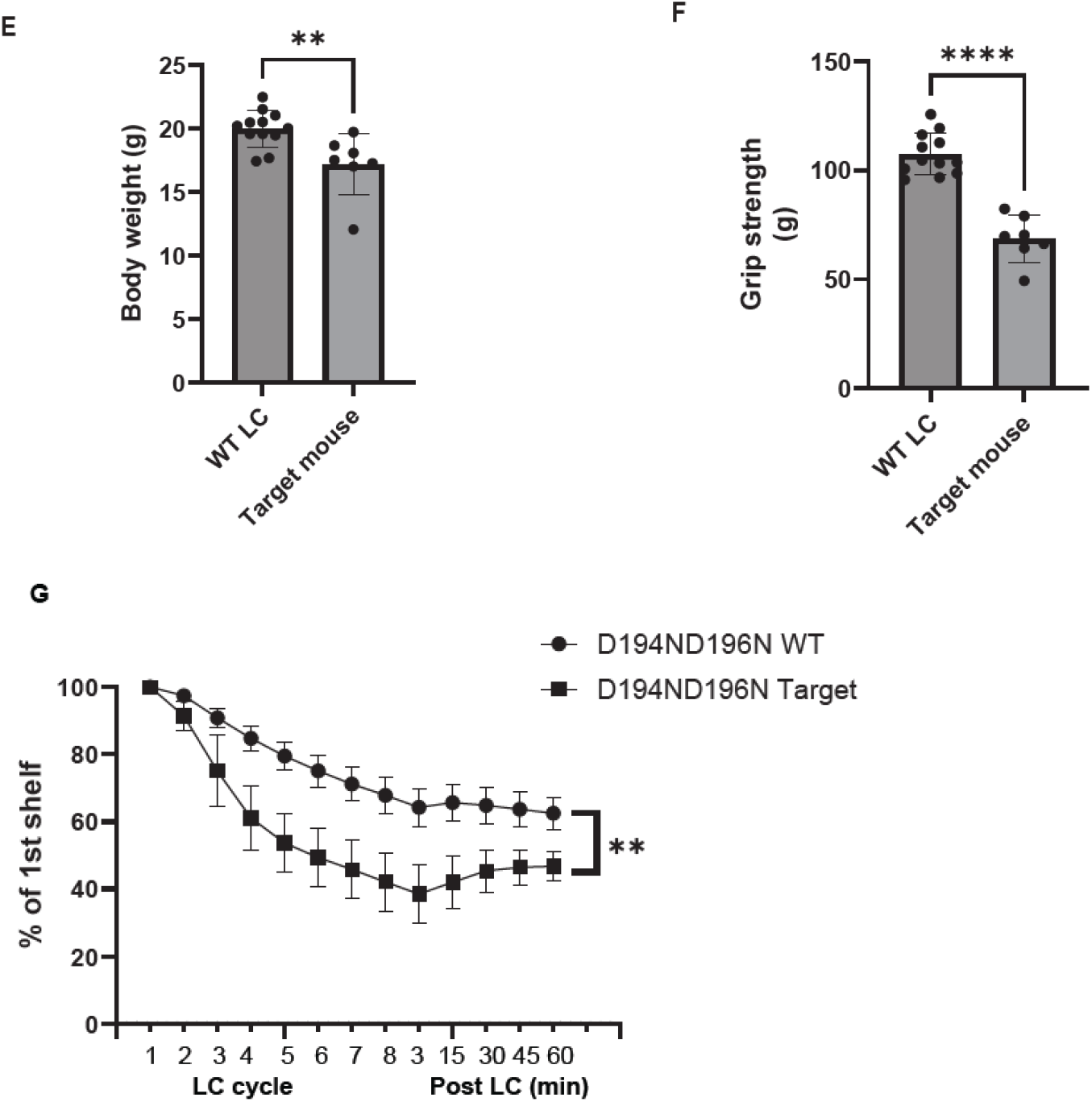
Xylose phosphatase activity of α-DGN is required for matriglycan extension. (**A**) D194N/D196N (DXD motif) mutations exhibited embryonic lethality in mice. (**B**) Strategy to create mice expressing muscle-specific DXD motif mutations (D194N/D196N) (target mice). (**C**) Immunoblot analysis of skeletal muscle to detect matriglycan (IIH6), core α & β DG (AF6868, the 100-200 kDa band represents α-DG and the discrete ∼ 42 kDa band is β-DG) and laminin overlay. Molecular weights are indicated on the left of each blot. (**D**) Clasping of hindlimbs in 7-week-old target mice (right side). (**E**) Body weight and (**F**) grip strength analysis of 7-week-old wild-type littermate control (WT LC) and target mice. Statistical significance determined by Student’s unpaired two-tailed t-test (**p-value= 0.005, ****p-value<0.0001). (**G**) Force recovery after lengthening contractions (LC) was measured. Statistical significance was determined by using unpaired two-tailed Student’s T test (*p value < 0.05, ****p value < 0.0001, **p value < 0.01, ns= not significant).

To test if the D194/D196 motif is required for matriglycan extension, we generated conditional mutants of D194N/D196N in mouse muscles using the Pax-Cre system (Fig. 5B) and called them “target mice”. Target mice showed a significant reduction in the molecular weight of matriglycan (∼90kD band) and DG (Fig. 5C). To test if the band observed in western blots that were probed with an anti-matriglycan antibody was indeed matriglycan, we performed a dual exoglycosidase digestion assay. The short matriglycan (∼90kD) was completely digested in the presence of exoglycosidases, confirming the band is matriglycan (fig. S6A and S6B).

Detailed investigation of muscle physiology in target mice revealed severe muscle pathology in the form of robust hindlimb clasping, reduced body weight, reduced grip strength, reduced absolute and specific tetanic force in muscle, reduced recovery from contraction-induced muscle injury and centrally nucleated muscle fibers (Fig. 5D-G and fig. S7A-G). These results indicate that the D194/D196 motif of α-DGN is likely the active site for xylose phosphatase activity, which is crucial for extension of matriglycan and therefore, muscle function.

## Discussion

Kinases and phosphatases act as a universal molecular switch in cell signaling mechanisms. Our investigation has revealed a unique mechanism wherein a part of a receptor (DG) acts as a phosphatase on its glycan modification to regulate it.

Matriglycan of different lengths is found on distinct sites on DG (*40*). FAM20B was identified in a dystroglycanopathy screen (*41*), however, its role in matriglycan formation was unknown. Our data suggests that FAM20B phosphorylation is required for initiation of matriglycan, but that, α-DGN-mediated dephosphorylation signals its extension, hence producing long or full-length matriglycan. Deletion of FAM20B is embryonic lethal in mice (*29*), which highlights the importance of xylose phosphorylation in matriglycan and glycosaminoglycan (GAG) synthesis. A recent mass spectrometry study of matriglycan identified an extra modification of approximately 80Da on the matriglycan precursor of some glycoforms, which matches the mass of a phosphate or a sulfate group (*40*). Our results suggest that this extra modification is from the xylose phosphate made by FAM20B. In addition to phosphates being an extremely labile modification, FAM20B-mediated xylose phosphorylation is also transient, which makes it challenging to be detected by mass spectrometry. Regardless, our work provides evidence that FAM20B xylose phosphorylation is required to facilitate initiation of matriglycan synthesis, and that α-DGN removes the phosphate to facilitate elongation.

Our data suggests that the removal of the phosphate by α-DGN is not immediate. The enzyme activity of α-DGN was highest with the phosphorylated primer disaccharide, indicating it may be the preferred substrate. However, α-DGN’s activity is much slower than FAM20B, suggesting that the slow activity of α-DGN gives LARGE1 enough time to initiate matriglycan synthesis. Our work shows that α-DGN can still dephosphorylate xylose after LARGE1 has added a xylose and/or a glucuronic acid (phosphorylated tri and tetra-saccharides), but at a reduced rate. This suggests that variation exists with regard to which step α-DGN dephosphorylates the xylose. Similarly, in case of GAG synthesis it is unclear at what stage the dephosphorylation of linker xylose occurs (*35*). Future studies are needed to fully understand the dynamics of this mechanism.

α-DGN is an autonomous globular domain (*42*) with two distinct subdomains (*43*). α-DGN’s S6 domain has an antiparallel beta-sheet with alpha-helices on one side of the sheet, whereas HAD-like phosphatases have a Rossman fold (*44*). DXD motif in the S6 domain of α-DGN is situated at the end of the beta-strand 1, similar to HAD-like phosphatases (*44*). This indicates that α-DGN’s HAD-like phosphatase activity is most likely a case of convergent evolution.

Notably, α-DGN is cleaved by the proprotein convertase furin in the trans-Golgi network after the sequence RVRR (amino acids 309-312) and is then secreted (*26, 45*). Although cleavage of α-DGN has no effect on the function of DG itself, our data suggest that α-DGN is crucial in Golgi to produce long polymers of matriglycan. α-DGN acts as a potential serum biomarker for Duchenne muscular dystrophy (*46*) and has been detected in a wide variety of human bodily fluids (*47, 48*). α-DGN is also shown to protect against proliferation of SARS-CoV-2 (*49*) and Influenza A virus (*37*). One study also shows that recombinant α-DGN can promote neurite extension in PC12 cells (*50*). The importance of α-DGN is underscored by the fact that its deletion is also embryonically lethal in mice (*23*) and here we have shown that mutations in the potential active site of α-DGN (D194N/D196N) also result in embryonic lethality. Despite its widespread function, the mechanisms by which α-DGN acts remain unknown. Our findings suggest that α-DGN could serve as a HAD-like xylose phosphatase. Given that all cell surfaces are coated with glycans or xylose containing glycans, our study provides a mechanism that could explain α-DGN’s diverse functions.

## Acknowledgments

Viral Vectors were provided by the University of Iowa Viral Vector Core (http://www.medicine.uiowa.edu/vectorcore). Transgenic mice were generated at the University of Iowa Genome Editing Core Facility directed by William Paradee, PhD and supported in part by grants from the NIH and from the Roy J. and Lucille A. Carver College of Medicine. We thank Norma Sinclair, Rongbin Guan, and Joanne Schwarting for their technical expertise in generating transgenic mice. We thank Dr. Miguel F. Gonzales of the Campbell Laboratory for help with retro-orbital injections in newborn mice. We thank Sally J. Prouty of the Campbell Laboratory for help with the H&E staining of muscle sections. We thank Dr. Erhard Hohenester of Imperial College London for helpful comments on the manuscript. Finally, we thank Dr. Jennifer Y. Barr of the scientific editing and research communications core at the University of Iowa for critical reading of the manuscript.

## Author contributions

Conceptualization: KPC, IC

Methodology: IC, DV, LY, KPC

Investigation: IC, DV, LY, BAW

Funding acquisition: KPC

Supervision: KPC

Writing – original draft: IC

Writing – review & editing: IC, KPC

## Data and materials availability

All data are available in the main text or the supplementary materials.

## Supplementary Materials

Materials and Methods

Figs. S1 to S7

References (*51-58*)

