## Supplementary Information for "Xylose phosphatase activity of dystroglycan self-regulates its receptor function"

**The PDF file includes:**

Materials and Methods
Figs. S1 to S7
References (51-58)

### Materials and methods

#### Preparation of -MU glycans

**Separation and purification of the primer disaccharide generated by B4GAT1dTM (GlcA-beta1,4-Xyl-MU).**

A large-scale reaction was carried out to produce GlcA beta1-4 Xylose-MU (G4X-MU), using recombinant His-tagged B4GAT1dTM (R & D Systems, 6664-GT-050). B4GAT1dTM (4ug in 400ul or 0.23uM) was added to 10 mM of UDP-GlcA (Sigma, U6751-100MG) and 1mM Xylose-β−MU (Glycosynth, 44111) in 50 mM MOPS buffer pH 6.0, 10 mM $MgCl_2$, 10 mM $MnCl_2$, and incubated for 8 hours at 37°C with rotation. The sample was then run over a C18 column (10mM × 250 mm Supelcosil LC-18 column (Supelco, Cat# 58368)) with Buffer A (50 mM ammonium formate pH 4.0) and Buffer B (80% acetonitrile (Fisher, A996-4) in buffer A), using a 12% B isocratic run at 2.5 ml/min on a Shimadzu Prominence HPLC system. The elution of MU derivatives was monitored by fluorescence detection (325 nm for excitation, and 380 nm for emission). The product GlcA beta1-4 Xylose-MU, in the peak fractions (around 20 minutes) was collected and lyophilized. The dried sample was then brought up in Milli-Q water (500 µl). The product was quantitated using a standard curve for GlcA-β-MU (Sigma, Cat # M3633).

**Separation and purification of the trisaccharide (Xylose alpha1,3-GlcA-beta1,4-Xyl-MU).**

A large-scale reaction was carried out to produce Xylose alpha1-3-GlcA beta1-4 Xylose-MU (XG4X-MU), using recombinant His-tagged like-acetylglucosaminyltransferase 1 deleted transmembrane domain (LARGE1dTM). LARGE1dTM (6 ug in 400 ul or 0.167 uM) purified using a metal-affinity resin as described previously (*14*) and was added to 10 mM of UDP-Xylose (Complex Carbohydrate Research Center- 10mg) and 1 mM GlcA-beta1,4-Xyl-MU in 50 mM MOPS buffer pH 6.0, 10 mM $MgCl_2$, 10 mM $MnCl_2$, and incubated for 8 hours at 37°C with rotation. The sample was then run over a C18 column (10mM × 250 mm Supelcosil LC-18 column (Supelco, Cat# 58368)) with Buffer A (50 mM ammonium formate pH 4.0) and Buffer B (80% acetonitrile in buffer A), using a 12% B isocratic run at 2.5 ml/min on a Shimadzu Prominence HPLC system. The elution of MU derivatives was monitored by fluorescence detection (325 nm for excitation, and 380 nm for emission). The product peak Xylose alpha1-3-GlcA beta1-4 Xylose-MU was collected and lyophilized. The dried sample was then brought up in Milli-Q water (500 µl). The product was quantitated using a standard curve for GlcA-β-MU.

**Separation and purification of the tetrasaccharide (GlcA-beta1,3-Xylose alpha1,3-GlcA-beta1,4-Xyl-MU).**

A large-scale reaction was carried out to produce GlcA beta1,3-xylose alpha1-3-GlcA beta1-4 Xylose-MU (GXG4X-MU), using recombinant His-tagged LARGE1dTM. LARGE1dTM (6 ug in 400 ul or 0.167 uM) was purified using a metal-affinity resin as described previously (*14*) and was added to 10 mM of UDP-GlcA (Complex Carbohydrate Research Center- 10 mg) and 1 mM of xylose alpha1,3-GlcA-beta1,4-Xyl-MU in 50 mM MOPS buffer pH 6.0, 10 mM $MgCl_2$, 10 mM $MnCl_2$, and incubated for 8 hours at 37°C with rotation. The sample was then run over a C18 column (10mM × 250 mm Supelcosil LC-18 column (Supelco, Cat# 58368)) with Buffer A (50 mM ammonium formate pH 4.0) and Buffer B (80% acetonitrile in buffer A), using a 12% B isocratic run at 2.5 ml/min on a Shimadzu Prominence HPLC system. The elution of MU derivatives was monitored by fluorescence detection (325 nm for excitation, and 380 nm for emission). The product peak GlcA beta1,3-xylose alpha1-3-GlcA beta1-4 Xylose-MU was collected and lyophilized. The dried sample was then brought up in Milli-Q water (500 µl). The product was quantitated using a standard curve for GlcA-β-MU.

**Separation and purification of the phosphorylated primer disaccharide generated by FAM20B (GlcA-beta1,4-Xyl-PO4-MU).**

A large-scale reaction was carried out to produce GlcA-beta1,4-Xyl-PO4-MU (G4Xp-MU), using recombinant His-tagged FAM20B (R & D Systems, 8427-FM-050). FAM20B (4 ug in 400 ul or 0.23 uM) was added to 1 mM of ATP (Sigma, A2383-1G) and 0.5 mM GlcA beta1-4 Xylose-MU, in 50 mM HEPES buffer pH 7.5, 10 mM $MnCl_2$, and incubated for 8 hours at 37°C with rotation. The sample was then run over a C18 column (10mM × 250 mm Supelcosil LC-18 column (Supelco, Cat# 58368)) with Buffer A (50 mM ammonium formate pH 4.0) and Buffer B (80% acetonitrile in buffer A), using a 12% B isocratic run at 2.5 ml/min on a Shimadzu Prominence HPLC system. The elution of MU derivatives was monitored by fluorescence detection (325 nm for excitation, and 380 nm for emission). The product GlcA beta1-4 Xylose-MU in the peak fractions (around 11 minutes) was collected and lyophilized. The dried sample was then brought up in Milli-Q water (500 µl). The product was quantitated using a standard curve for GlcA-β-MU.

**Separation and purification of the phosphorylated trisaccharide (Xylose alpha1,3-GlcA-beta1,4-Xyl-PO4-MU).**

A large-scale reaction was carried out to produce Xylose alpha1-3-GlcA beta1-4 Xylose-PO4-MU (XG4Xp-MU), using recombinant His-tagged LARGE1dTM. LARGE1dTM (6 ug in 400 ul or 0.167 uM) was purified using a metal-affinity resin as described previously (*14*) and was added to 10 mM of UDP-Xylose (Complex Carbohydrate Research Center- 10 mg) and 1mM GlcA-beta1,4-Xyl-PO4-MU in 50 mM MOPS buffer pH 6.0, 10 mM $MgCl_2$, 10 mM $MnCl_2$, and incubated for 8 hours at 37°C with rotation. The sample was then run over a C18 column (10 mM × 250 mm Supelcosil LC-18 column (Supelco, Cat# 58368)) with Buffer A (50 mM ammonium formate pH 4.0) and Buffer B (80% acetonitrile in buffer A), using a 12% B isocratic run at 2.5 ml/min on a Shimadzu Prominence HPLC system. The elution of MU derivatives was monitored by fluorescence detection (325 nm for excitation, and 380 nm for emission). The product peak Xylose alpha1-3-GlcA beta1-4 Xylose-PO4-MU was collected and lyophilized. The dried sample was brought up in Milli-Q water (500 µl). The product was quantitated using a standard curve for GlcA-β-MU.

**Separation and purification of the phosphorylated tetrasaccharide (GlcA-beta1,3-Xylose alpha1,3-GlcA-beta1,4-Xyl-PO4-MU).**

A large-scale reaction was carried out to produce GlcA beta1,3-xylose alpha1-3-GlcA beta1-4 Xylose-PO4-MU (GXG4Xp-MU), using recombinant His-tagged LARGE1dTM. LARGE1dTM (6 ug in 400 ul or 0.167 uM) was purified using a metal-affinity resin as described previously (*14*) and was added to 10 mM of UDP-GlcA (Complex Carbohydrate Research Center- 10 mg) and 1 mM xylose alpha1,3-GlcA-beta1,4-Xyl-MU in 50 mM MOPS buffer pH 6.0, 10 mM $MgCl_2$, 10 mM $MnCl_2$, and incubated for 8 hours at 37°C with rotation. The sample was then run over a C18 column (10 mM × 250 mm Supelcosil LC-18 column (Supelco, Cat# 58368)) with Buffer A (50 mM ammonium formate pH 4.0) and Buffer B (80% acetonitrile in buffer A), using a 12% B isocratic run at 2.5 ml/min on a Shimadzu Prominence HPLC system. The elution of MU derivatives was monitored by fluorescence detection (325 nm for excitation, and 380 nm for emission). The product peak GlcA beta1,3-xylose alpha1-3-GlcA beta1-4 Xylose-PO4-MU was collected and lyophilized. The dried sample was then brought up in Milli-Q water (500 µl). The product was quantitated using a standard curve for GlcA-β-MU.

**Separation and purification of the trisaccharide (GGM-MU) and the phosphorylated trisaccharide (GGMp-MU).**

A large-scale reaction was carried out to produce GalNAc- beta1-3 GlcNAc beta1-4 Mannose-PO4-MU, using recombinant His-tagged B3GALNT2 (*38*) and the sugar GlcNAc beta1-4 Mannose-4- methylumbelliferone (MU) (Sussex Research, product # ES631010). B3GALNT2dTM (5ug in 400ul or 0.23uM) was added to 10 mM of UDP-GalNAc (Sigma, U5252-100MG), 1 mM GlcNAc beta1-4 Mannose-4- methylumbelliferone (MU), in 50 mM MOPS buffer pH 6.0, 10 mM $MgCl_2$, 10 mM $MnCl_2$, and incubated for 8 hours at 37°C with rotation. The sample was then run over a C18 column (10mM × 250 mm Supelcosil LC-18 column (Supelco, Cat# 58368)) with Buffer A (50 mM ammonium formate pH 4.0) and Buffer B (80% acetonitrile in buffer A), using a 16% B isocratic run at 2.5 ml/min on a Shimadzu Prominence HPLC system. The elution of MU derivatives was monitored by fluorescence detection (325 nm for excitation, and 380 nm for emission). The product GalNAc- beta1-3 GlcNAc beta1-4 Mannose-MU was collected in the peak fractions (around 20 minutes) and lyophilized. The dried sample was then brought up in Milli-Q water (500 µl). The product was quantitated using a standard curve for GlcA-β-MU. The second step is to add phosphate to the trisaccharide GalNAc- beta1-3 GlcNAc beta1-4 Mannose-PO4-MU. A large-scale reaction was carried out to produce GalNAc- B1-3 GlcNAc B1-4 Mannose-PO4-MU, using recombinant His-tagged POMK. POMK (5 ug in 400 ul or 0.23 uM) was added to 10 mM of ATP (Sigma, A2383-100MG), 1 mM 4-Methylumbelliferone alpha-GlcNAc B 1-4 Mannose, in 50 mM MOPS buffer pH 6.5, 10 mM $MgCl_2$, 10 mM $MnCl_2$, and incubated for 8 hours at 37°C with rotation. The sample was then run over a C18 column (10 mM × 250 mm Supelcosil LC-18 column (Supelco, Cat# 58368)) with Buffer A (50 mM ammonium formate pH 4.0) and Buffer B (80% acetonitrile in buffer A), using a 16% B isocratic run at 2.5 ml/min on a Shimadzu Prominence HPLC system. The elution of MU derivatives was monitored by fluorescence detection (325 nm for excitation, and 380 nm for emission). The product GalNAc- B1-3 GlcNAc B1-4 Mannose-PO4-MU was collected in the peak fractions (around 12 minutes) and lyophilized. The dried sample was then brought up in Milli-Q water (500 µl). The product was quantitated using a standard curve for GlcA-β-MU.

**FAM20B assay**

FAM20B protein (R&D Systems, 8427-FM-050) was reconstituted as a 500 µg/ml stock in 25 mM Tris-HCL. To assay FAM20B activity with G4-MU (25 µM) and G5-MU (20 µM) [LARGE1 synthesized glycans, described previously (*30*)] 10 µg/ml of FAM20B was used with 0.2 mM ATP and 10 mM $MnCl_2$ in a 50 mM HEPES pH 7.4 buffer in a total reaction volume of 150 µl. Reaction along with control (No FAM20B) was incubated at 30°C for 16 hours with 300 rpm shaking. 20 ul reaction and control samples were taken at 1hr, 3hr, 5hr and 16hr and ran on C-18 HPLC with Buffer A (50 mM ammonium formate pH 4.0) and Buffer B (80% acetonitrile in Buffer A), using a 9% B isocratic run at 2.5 ml/min.

For the FAM20B assay with G4X-MU (prepared as described above), 20 µg/ml of FAM20B was incubated with 25 µM G4X-MU with or without 0.2 mM ATP, 10 mM $MnCl_2$, 50 mM HEPES pH 7.4 in a total reaction volume of 200 µl. Reaction and control (without FAM20B) were incubated at 37°C with 300 rpm shaking. Samples were taken at 2hr, 4hr, 6hr and 16hr and separated on C18 HPLC with Buffer A (50 mM ammonium formate pH 4.0) and Buffer B (80% acetonitrile in Buffer A), using a 12% B isocratic run at 2.5 ml/min.

For assay with Xylose-beta-MU, 25 µM of glycan was incubated with 20 µg/ml of FAM20B, 0.2 mM ATP, 10 mM $MnCl_2$, 50 mM HEPES pH 7.4 in a total reaction volume of 200 µl and kept at 37°C with 300 rpm shaking along with a control reaction without FAM20B. Samples were taken at 2hr, 4hr, 6hr and 16hr and ran on C18 HPLC with Buffer A (50 mM ammonium formate pH 4.0) and Buffer B (80% acetonitrile in Buffer A), using a 12% B isocratic run at 2.5 ml/min.

For assays with XG4X-MU and GXG4X-MU, 7 µM of each glycan was incubated with 10 µg/ml of FAM20B, 0.2 mM ATP, 10 mM $MnCl_2$, 50 mM HEPES pH 7.4 in a total reaction volume of 200 µl and kept at 37°C with 300 rpm shaking along with a control reaction without FAM20B. Samples were taken at 2hr, 4hr, 6hr and 16hr and ran on C18 HPLC with Buffer A (50 mM ammonium formate pH 4.0) and Buffer B (80% acetonitrile in Buffer A), using a 12% B isocratic run at 2.5 ml/min.

For FAM20B assay using HAP cell lysates, WT C631 and FAM20B KO cell lines (Horizon Discovery, catalog # HZGHC006909c018) were grown in 150 mm plates in IMDM media (with 10% Fetal Bovine Serum and 1% Penicillin/Streptomycin) and collected upon reaching 100% confluency using a cell scraper. Cells were spun at 14000 rpm at 4°C for 10 minutes and resuspended in 500 µl homogenization buffer (50 mM Tris pH 7.6, 150 mM NaCl, 1% TX-100, 10 mM EDTA and all protease inhibitors). Cells were vortexed 3 times for 1 minute each and rotated in a cold room for 1 hour to complete cell lysis. Cells were then spun down at 14000 rpm for 10 minutes at 4$^o$C. Supernatant was collected as the cell lysate. 15 µl of each WT and FAM20B HAP cell lysate was incubated with 25 µM of G4X-MU in 50 mM HEPES pH 7.4 buffer, 0.2 mM ATP and 10 mM $MnCl_2$ in a total reaction volume of 200 µl. Reactions were incubated at 37$^o$C at 300 rpm and samples were collected at 3hr and 16hr and ran on C18 HPLC with Buffer A (50 mM ammonium formate pH 4.0) and Buffer B (80% acetonitrile in Buffer A), using a 12% B isocratic run at 2.5 ml/min.

**FAM20C Assay**

Recombinant human FAM20C protein was purchased from R&D Systems (9265-FM-050) and reconstituted as a 500 µg/ml stock in 25 mM Tris-HCL. G4X-MU (25 µM) was incubated with FAM20C (20 µg/ml) in the presence of 0.2 mM ATP, 10 mM $MnCl_2$ and 50 mM HEPES buffer pH 7.5 and incubated at 37$^o$C with 300 rpm shaking for 17 hours along with a control reaction without FAM20C. Samples were taken at 2hr, 6hr and 17hr. and ran on C18 HPLC column with Buffer A (50 mM ammonium formate pH 4.0) and Buffer B (80% acetonitrile in Buffer A), using a 12% B isocratic run at 2.5 ml/min.

**FAM198A assay**

Recombinant human FAM198A (also called GASK1A) was purchased from Origene (TP329292) and reconstituted at 150 µg/ml. G4X-MU (1µM) was incubated with FAM198A (5 µg/ml) in the presence of 0.2 mM ATP, 10 mM $MnCl_2$, 10 mM $MgCl_2$ in 50 mM HEPES pH7.4 buffer (total reaction volume 200 µl) and incubated for 16 hours at 37$^o$C with 300 rpm shaking along with a control reaction without FAM198A. Samples were taken at 2hr, 4hr, 6hr and 16hr and ran on C18 HPLC column with Buffer A (50 mM ammonium formate pH 4.0) and Buffer B (80% acetonitrile in Buffer A), using a 12% B isocratic run at 2.5 ml/min.

**FJX-1 (Four jointed 1) assay**

Recombinant human FJX-1 protein was purchased from R&D Systems (H00024147-P01-2ug) and reconstituted at 40 µg/ml. G4X-MU (500 nM) was incubated with FJX-1 (10 µg/ml) in the presence of 0.2 mM ATP, 10 mM $MnCl_2$, 10 mM $MgCl_2$ in 50 mM HEPES pH 7.4 buffer (total reaction volume 100 µl) and incubated for 16 hours at 37$^o$C with 300 rpm shaking along with a control reaction without FJX-1. Samples were taken at 16 hr and ran on a C18 HPLC column with Buffer A (50 mM ammonium formate pH 4.0) and Buffer B (80% acetonitrile in Buffer A), using a 12% B isocratic run at 2.5 ml/min.

**LARGE1 assay**

Matriglycan was synthesized using either G4X-MU (20 µM) or G4Xp-MU (20 µM) as starting substrate. LARGE1dTM (0.5 µM, prep 6) was incubated with the disaccharides in the presence of 5 mM UDP-Xylose, 5 mM UDP-Glucuronic acid, 10 mM $MnCl_2$, 10 mM $MgCl_2$ and 50 mM MOPS buffer pH 6.0 in a 250 µl reaction volume. The reactions were incubated at 37$^o$C with 300 rpm shaking for 90 minutes. Reactions were stopped by heating at 99°C for 5 minutes. Products were separated using a SAX (strong anion exchange) column with a NaCl gradient and quantified based on the area under the peak.

**Production and purification of N-terminal domain of dystroglycan (α-DGN)**

Mammalian cells stably expressing His-tagged rabbit α-DGN were used as described previously (*37*). Briefly, HEK293 cells were transfected with the pcDNA3.1 mammalian expression vector (Thermo Fisher Scientific, Waltham, MA) containing the complete N-terminal sequence of rabbit α-DGN with the His-tag inserted directly after the signal peptide sequence using FuGene 6 Transfection Reagent (Promega, Madison, WI). Cells stably expressing His-tagged rabbit α-DGN were cultured in DMEM supplemented with G418 (Thermo Fisher Scientific, Waltham, MA). HEK293 cells stably expressing His-tagged rabbit α-DGN were adapted to serum-free medium, 293 SFM II (Thermo Fisher Scientific, Waltham, MA), and cultivated in CELLine bioreactors (CL1000; Argos Technologies, Vernon Hills, IL). His-tagged rabbit α-DGN secreted into the culture medium was purified using the Talon metal-affinity resin (Takara Bio USA Inc., Mountain View, CA) according to the manufacturer's instructions. The purity of the protein was confirmed by SDS-PAGE and Coomassie Brilliant Blue staining.

Further, to remove high molecular weight proteins and obtain a pure α-DGN, we subjected the eluates from Talon metal affinity purification to size exclusion chromatography (SEC). Relevant Talon eluates were combined, concentrated and buffer exchanged with 1X PBS pH 7.4 (to remove imidazole) in Amicon centrifugal filters (MWCO 10,000Da). HiLoad 16/600 Superdex 200 pg column was preequilibrated with 20 mM HEPES pH 7.4 buffer at 0.1 ml/min flow rate. Approximately 1 ml of Talon eluate was passed through the preequilibrated column at a flow rate of 1 ml/min. Fractions were collected for up to 180 minutes. Relevant fractions were run on an SDS-PAGE gel and stained with Coomassie Brilliant Blue to confirm purity. Fractions showing pure α-DGN were collected, combined and concentrated in Amicon centrifugal filters (MWCO 10,000 Da). Pure α-DGN was used in enzyme assays. The concentration of α-DGN was determined using A280 absorbance measurements by NanoDrop.

**DGN assays**

Pure α-DGN (40 μg/ml, 1$^{st}$ prep) was incubated with G4Xp-MU (0.5 μM or 2 μM or 7 μM) in 50 mM HEPES pH 7.4 or 50 mM MES pH 6 buffers with or without 10 mM $MgCl_2$, with or without 10 mM $MnCl_2$ and with or without 20 mM EDTA, wherever indicated. Reactions along with their controls were incubated at 37°C with 300 rpm shaking for 16 hours. Samples were taken at 2hr, 4hr and 16hr and ran on a C18 HPLC column with Buffer A (50 mM ammonium formate pH 4.0) and Buffer B (80% acetonitrile in Buffer A), using a 12% B isocratic run at 2.5 ml/min.

Pure α-DGN (47 μg/ml, 2$^{nd}$ prep) was incubated with G4Xp-MU (2 μM), GGMp-MU (2 μM), XGX4p-MU (2 μM) and GXG4Xp-MU (2 μM) with 10 mM $MgCl_2$ and 50 mM MES pH 6.0 buffer at 37°C with 300 rpm shaking for 16 hours along with controls without α-DGN. Samples were taken at 2hr, 4hr, 6hr and 16hr and ran on a C18 HPLC column with Buffer A (50 mM ammonium formate pH 4.0) and Buffer B (80% acetonitrile in Buffer A), using a 12% B isocratic run at 2.5 ml/min.

**NMR spectroscopy**

All NMR spectra were acquired at 25°C on a Bruker Avance II 800 MHz spectrometer equipped with a TCI cryoprobe, or a Bruker Avance NEO 600 MHz spectrometer equipped with a QCI-P cryoprobe. For structure determination of the phosphorylated product (0.984 mM) of FAM20B enzymatic reaction with the substrate G4X-MU, one-dimensional (1D) $^{1}H$ spectra, $^{1}H$ homonuclear two-dimensional (2D) DQF-COSY, TOCSY, and ROESY spectra, and $^{1}H/^{13}C$ 2D heteronuclear HMQC and H2BC spectra were collected at the 800 MHz NMR spectrometer. The $^{31}P$ 1D and $^{31}P/^{1}H$ COSY spectrum (*51*) were acquired at the 600 MHz NMR spectrometer. Glycan binding studies by LARGE1 were all carried out at the 800 MHz NMR spectrometer. 1D $^{1}H$ NMR spectra of the glycan (G4X-MU or G4Xp-MU) in the absence and presence of various concentrations of LARGE1 were acquired using a 50 ms $T_2$ filter consisting of a train of spin-lock pulses to eliminate the broad resonances from the protein (*52*). LARGE1 titrations with glycans were performed in a buffer containing 20 mM HEPES and 150 mM NaCl, pH 7.4 in 100% $D_2O$. The $^{13}C$ and $^{1}H$ resonances of the LARGE acceptor primer glycan were reported previously (*13*). The $^{1}H$ chemical shifts are referenced to 2,2-dimethyl-2-silapentane-5-sulfonate. The collected data were processed using NMRPipe (*53*) and analyzed using NMRView (*54*). Glycan binding affinity to LARGE1 was determined using glycan-observed NMR experiments as described previously (*30*). For the resolved anomeric Xyl H1 peak, the bound fraction was calculated by measuring the difference in the peak intensity in the absence (free form) and presence (bound form) of LARGE1, and then dividing by the peak intensity of the free form. To obtain dissociation constants, the data were fitted to the standard quadratic equation using GraphPad Prism (GraphPad Software). The standard deviation from data fitting is reported.

**Animals**

All mice were maintained in a barrier-free, specific pathogen-free grade facility and had access to normal chow and water ad libitum. All animals were manipulated in biosafety cabinets and change stations using aseptic procedures. The mice were maintained in a climate-controlled environment at 25°C on a 12/12hr light/dark cycle. Animal care, ethical usage, and procedures were approved and performed in accordance with the standards set forth by the National Institutes of Health and the University of Iowa Animal Care and Use Committee (IACUC). Mouse lines used in the study that have been previously described are: $Dag1^{flox}$ (JAX#009652) (*10*) and $Pax7^{cre}$ (JAX# 010530) (*58*). Littermate controls were employed whenever possible. The number of animals required and the number of replicants performed were based on experience with standard deviations of the given techniques. The number of mice used is provided in each figure legend. Male and female mice were analyzed separately in physiological assays. No outliers were encountered, and no mice were excluded.

**Generation of mouse lines**

The $Dag1^{D121ND123N}$, $Dag1^{D196N}$ and $Dag1^{D194ND196N}$ mouse lines were developed by the Genome Editing Facility at the University of Iowa.

To generate the D121N/D123N mutation, point mutations were introduced (GAC>AAC and GAT>AAT at positions 9983 and 9989 of the genomic DNA sequence) using CRISPR. The Cas-9 protein, single guide RNA and repair donor DNA (ssODN) were injected into the mouse zygote pronucleus. Guide RNA sequence: 5' ACACCTTTATCAGTGTCAAG 3'. The founder mouse was identified through genomic DNA sequencing using the following primers: Forward: 5' CCTCAGGTTAGAGCTTGGTTTAT 3' Reverse: 5'CTCATTGTGGTCTTCAGGGTAG 3'. The founder mouse was then crossed to a C57BL6/J mouse to obtain N1 pups. N1 pups were sequenced again to test germline transfer of the mutations. The N1 pups that successfully carried the mutations were crossed again to a C57BL6/J mouse to obtain N2 generation. Heterozygous progenies were crossed again to obtain homozygotes (F1).

To generate D194N/D196N mutation, point mutations were introduced (GAT>AAT and GAC>AAT at positions 10,202, 10,208 and 10,210 of the genomic DNA sequence) using CRISPR. The Cas-9 protein, single guide RNA and repair donor DNA (ssODN) were injected into the mouse zygote pronucleus. Guide RNA sequence: 5' GCTTTGGGGTCATCTTGGTG 3'. The founder mouse was identified through genomic DNA sequencing using the following primers: Forward: 5' GGGACCCACACAGTCATATTT 3' Reverse: 5' GTTCAAGGAGCAGCCTAGTT 3'. The founder mouse was then crossed to a C57BL6/J mouse to obtain N1 pups. N1 pups were sequenced again to test germline transfer of the mutations. The N1 pups that successfully carried the mutations were crossed again to a C57BL6/J mouse to obtain the N2 generation. Heterozygous progenies were crossed again to obtain homozygotes (F1). Since no homozygous mice could be obtained for this line, F1 heterozygotes were crossed to heterozygous $Pax7^{cre}$; $Dag1^{flox}$ mice to obtain target mice.

To generate the D196N mutation, a point mutation was introduced (GAC>AAC at position 10,208 of the genomic DNA sequence) using CRISPR. The Cas-9 protein, single guide RNA and repair donor DNA (ssODN) were injected into the mouse zygote pronucleus. Guide RNA sequence: 5' GCTTTGGGGTCATCTTGGTG 3'. The founder mouse was identified through genomic DNA sequencing using the following primers: Forward: 5' GGGACCCACACAGTCATATTT 3' Reverse: 5' GTTCAAGGAGCAGCCTAGTT 3'. The founder mouse was then crossed to a C57BL6/J mouse to obtain N1 pups. N1 pups were sequenced again to test germline transfer of the mutations. The N1 pups that successfully carried the mutations were crossed again to a C57BL6/J mouse to obtain the N2 generation. Heterozygous progenies were crossed again to obtain homozygotes (F1).

**Generation of FAM20B knockdown mice**

Adenovirus5-U6-m-FAM20B-shRNA was purchased from Vector Biolabs (SKU shADV-259046) and amplified by the Viral Vector Core at the University of Iowa to obtain a high titer of 1 x $10^{11}$ pfu/ml. The virus was further sequenced to confirm successful amplification. Nine two-day-old C57BL6/J mice were injected with 10µl of the high-titer virus through the retro-orbital sinus route. Mice that survived (4) were euthanized after 3 weeks of injections and tissues were harvested from them and their un-injected littermates.

**Forelimb grip strength test**

Forelimb grip strength was measured at 3 months using previously published methods (*21, 37*). A mouse grip strength meter (Columbus Instruments, Columbus, OH, USA) was mounted horizontally, with a non-flexible grid connected to the force transducer. The mouse was allowed to grasp the grid with its two front paws and then pulled away from the grid by its tail until the grip was broken. This was done three times over five trials, with a 1 min break between each trial. The gram force was recorded per pull, and any pull where only one front limb or any hind limbs were used was discarded. If the mouse turned, the pull was also discarded. After 15 pulls (five sets of three pulls), the mean of the three highest pulls of the 15 was calculated and reported. Statistics were calculated using GraphPad Prism 8 software. One-way ANOVA and two-sided Student's t-test were used accordingly. Differences were considered significant at a p-value less than 0.05. Graph images were also created using GraphPad Prism and the data in the present study are shown as the means ± SD unless otherwise indicated.

**Body weight measurements**

Mice were weighed as previously described (*21, 37*). Weights were measured after testing grip strength using a Scout SPX222 scale (OHAUS Corporation, Parsippany, NJ, USA), and the tester was blinded to genotype. Statistics were calculated using GraphPad Prism 8 software and one-way ANOVA and two-tailed Student's t-test was used accordingly. Differences were considered significant at a p-value less than 0.05. Graph images were also created using GraphPad Prism and the data in the present study are shown as the means ± SD unless otherwise indicated.

**Measurement of in vitro muscle function**

To compare the contractile properties of muscles, EDL muscles were surgically removed as described previously (*55*). The muscle was immediately placed in a bath containing a buffered physiological salt solution (composition in mM: NaCl, 137; KCl, 5; CaCl2, 2; $MgSO_4$, 1; $NaH_2PO_4$, 1; $NaHCO_3$, 24; glucose, 11). The bath was maintained at 25°C, and the solution was bubbled with 95% $O_2$ and 5% $CO_2$ to stabilize pH at 7.4. The proximal tendon was clamped to a post and the distal tendon was tied to a dual-mode servomotor (Model 305C; Aurora Scientific, Aurora, ON, Canada). Optimal current and optimal whole muscle length ($L_0$) were determined by monitoring isometric twitch force. Optimal frequency and maximum isometric tetanic force

($F_0$) were also determined. The muscle was then subjected to an EC protocol consisting of eight ECs at 3 min intervals. A fiber length $L_f$-to- $L_0$ ratio of 0.45 was used to calculate $L_f$. Each EC consisted of an initial 100 ms isometric contraction at optimal frequency immediately followed by a stretch of Lo to 30% of $L_f$ beyond $L_o$ at a velocity of 1 $L_f$/s at optimal frequency. The muscle was then passively returned to $L_o$ at the same velocity. At 3, 15, 30, 45, and 60 min after the EC protocol, isometric tetanic force was measured. After the analysis of the contractile properties, the muscle was weighed. The CSA of muscle was determined by dividing the muscle mass by the product of $L_f$ and the density of mammalian skeletal muscle (1.06 g/cm$^3$). The specific force was determined by dividing $F_o$ by the CSA (kN/mm$^2$). Eleven- to 16-week-old male and female mice were used, and right EDL muscles from each mouse were employed, with three to four muscles used for each analysis. Each data point represents an individual EDL. Statistics were calculated using GraphPad Prism 8 software and one-way ANOVA and two-tailed Student's t test were used accordingly. Differences were considered significant at a p-value less than 0.05.

**Tissue biochemical analysis**

One gram of mouse skeletal muscle or one brain or one heart from individual mouse was turned to powder in liquid nitrogen using a mortar and pestle, then homogenized with polytron (Kinematica, PT10-35) three times for 10 s at power 4–5 in 6 mL of TBS (50 mM Tris & 150 mM NaCl) with 1% TX-100 and 10 mM EDTA, and protease inhibitors (at final concentrations: 0.1 mM phenylmethylsulfonylfluoride (PMSF)), 0.75 mM benzamidine,0.5 µg/ml leupeptin (Sigma/Millipore), 0.6 µg/ml pepstatin A (Millipore), 0.5 µg/ml aprotinin (Sigma-Aldrich), 5 µM Calpain I inhibitor and Calpeptin. The samples were incubated in a cold room for 1hr with rotation. The samples were centrifuged in a Beckman Coulter Avanti J-E centrifuge for 30 minutes at 20,000 *g*, 4°C. The supernatant was combined with WGA slurry (Vector Laboratories, AL-1023) at 500 µL per gram of starting muscle and rotated at 4°C overnight. The WGA beads were washed using 5 ml of WGA wash buffer 4x for 5 minutes at 1500 *g* with 0.1% Tx/TBS, plus protease inhibitors. After the final wash, the WGA beads were mixed with 5x Laemmli Sample Buffer (LSB) at a 1:1 ratio and heated at 99°C for 10 minutes. Samples were loaded (beads and LSB) in a 3–15% gradient gel. The proteins were transferred to PVDF-FL membranes (Millipore) as previously published (*4,19*). EDTA (10 mM) was used in the homogenization to extract α-DG containing matriglycan more efficiently.

**Immunoblotting and ligand overlay**

The mouse monoclonal antibody against matriglycan on α-DG (IIH6, Developmental Studies Hybridoma Bank, University of Iowa; RRID: AB_2617216) was characterized previously and used at 1:50 (*56*). The polyclonal antibody, AF6868 (R&D Systems, Minneapolis, MN, USA; RRID: AB_10891298), was used at a concentration of 1:200 for immunoblotting the core α-DG and β-DG proteins, and the secondary was a donkey anti-sheep (LI-COR Bioscience, Lincoln, NE, USA) used at 1:10,000 concentration. The monoclonal anti-FAM20B antibody (R&D,

Systems, Catalog #: MAB8427) was used at a 1:200 concentration to detect FAM20B protein in mouse and human cell lines. The secondary antibody used was goat anti-mouse $IgG_1$ at 1:3000 concentration. The Blots were developed with infrared (IR) dye-conjugated secondary antibodies and scanned using the Odyssey infrared imaging system (LI-COR Bioscience). Blot images were captured using the included Odyssey image-analysis software. Laminin overlay assays were performed as previously described (*4,19*). Immobilon-FL membranes were blocked in laminin-binding buffer (LBB: 10 mM triethanolamine, 140 mM NaCl, 1 mM $MgCl_2$, 1 mM $CaCl_2$, pH 7.6) containing 5% non-fat dried milk, followed by incubation with mouse Engelbreth-Holm-Swarm laminin (Thermo Fisher, Catalog#: 23017015) overnight at a concentration of 7.5 nM at 4°C in LBB containing 3% bovine serum albumin (BSA) and an additional 2 mM $CaCl_2$. Membranes were washed and incubated with anti-laminin antibody (L9393; Sigma-Aldrich 1:1000 dilution) followed by IRDye 800 CW dye-conjugated donkey anti-rabbit IgG (LI-COR, 926-32213) at 1:2500.

**Digestion of α-DG with exoglycosidases**

The digestion was performed as previously described (*57*). Briefly, β-Glucuronidase from *T. maritima* and α-xylosidase from *S. solfataricus* were cloned into pET-28a (+) vector between *Nhe*I/*Xho*I sites in frame with the N-terminal 6xHis tag. The plasmids (20 ng each) were chemically transformed into 50 µL BL21DE3 One Shot competent cells. One colony each was picked and inoculated in 20 mL LB (with kanamycin 50 µg/mL) overnight at 37°C. The next day, 10 mL of the overnight culture was inoculated into 1 L LB (with kanamycin 50 µg/mL). After reaching 0.6 OD at 600 nm, the cultures were induced with 1 mM IPTG and incubated at 16°C overnight. The next day, the cells were centrifuged at 5000. *g*, for 10 min at 4°C. Cell pellets were stored at –80°C until ready for purification. The pellets were dissolved in 20 mL homogenization buffer (50 mM Tris-Cl, 150 mM NaCl, 1% TX-100, and all protease inhibitors) per liter culture. The cells were stored again overnight in 50 mL Falcon tubes at –80°C for ice crystal formation. Cells were thawed the next day for purification. Nuclease (Pierce) was added at 1.25 kU and cells were sonicated at power level four-five for four times with 10 s intervals in between at 4°C. Cells were then centrifuged at 15,000 *g* for 20 minutes at 4°C. The supernatant was heat fractionated at 75°C for 10 min after which it was centrifuged at 15,000 *g* for 30 minutes at 4ºC. Meanwhile, a TALON Superflow metal affinity column was prepared by packing 3 mL of resin and equilibrating with wash buffer 1 (50 mM Tris-Cl, 100 mM NaCl, 0.1% TX-100, all PIs). All further purification steps were performed at 4°C. The extract was applied to the column three times, such that each time, the extract was incubated with the column for 15–30 min on gentle rocking platform. All flowthrough was saved. The column was washed three times with wash buffer 1. All washes were saved. The column was next washed with a high salt wash buffer (50 mM Tris-Cl, 500 mM NaCl, 0.1% TX-100, all PIs) to remove nonspecific interactions and the high salt wash was saved. Proteins were then eluted with elution buffer (50 mM Tris-Cl, 100 mM NaCl, 0.1% TX-100, and 300 mM Imidazole) in five fractions of 3 mL each. The relevant fractions (elutes 1 and 2) were pooled together, and buffer exchanged with 1XPBS pH

7.4 with 30 kDa concentrators (Amicon). One hundred µL was loaded on SDS-PAGE from all fractions and washes to visualize with Coomassie. WGA-enriched glycoproteins (elutes) were buffer exchanged with sodium acetate buffer pH 5.5 using 30 kDa concentrators and heated for 5 min in the presence of 10 mM β-mercaptoethanol at 99°C. All protease inhibitors were added after the mixture cooled down. Fifty µL of each enzyme was added per 250 µL of WGA-enriched and buffer-exchanged glycoproteins. The initial time point was aliquoted as $T_o$ and the rest was incubated at 75°C with 600 rpm shaking for 16hr.

**H&E analysis of skeletal muscle**

Histology and immunofluorescence of mouse skeletal muscle were performed as described previously (*19*). Mice were euthanized by cervical dislocation and directly after sacrifice, quadriceps muscles were isolated, embedded in OCT compound, and then snap-frozen in liquid nitrogen-cooled 2-methylbutane. Ten µm sections were cut with a cryostat (Leica CM3050S Research Cryostat; Amsterdam, The Netherlands) and H&E stained using conventional methods. Whole digital images of H&E-stained sections were taken by a VS120-S5-FL Olympus slide scanner microscope (Olympus Corporation, Tokyo, Japan).

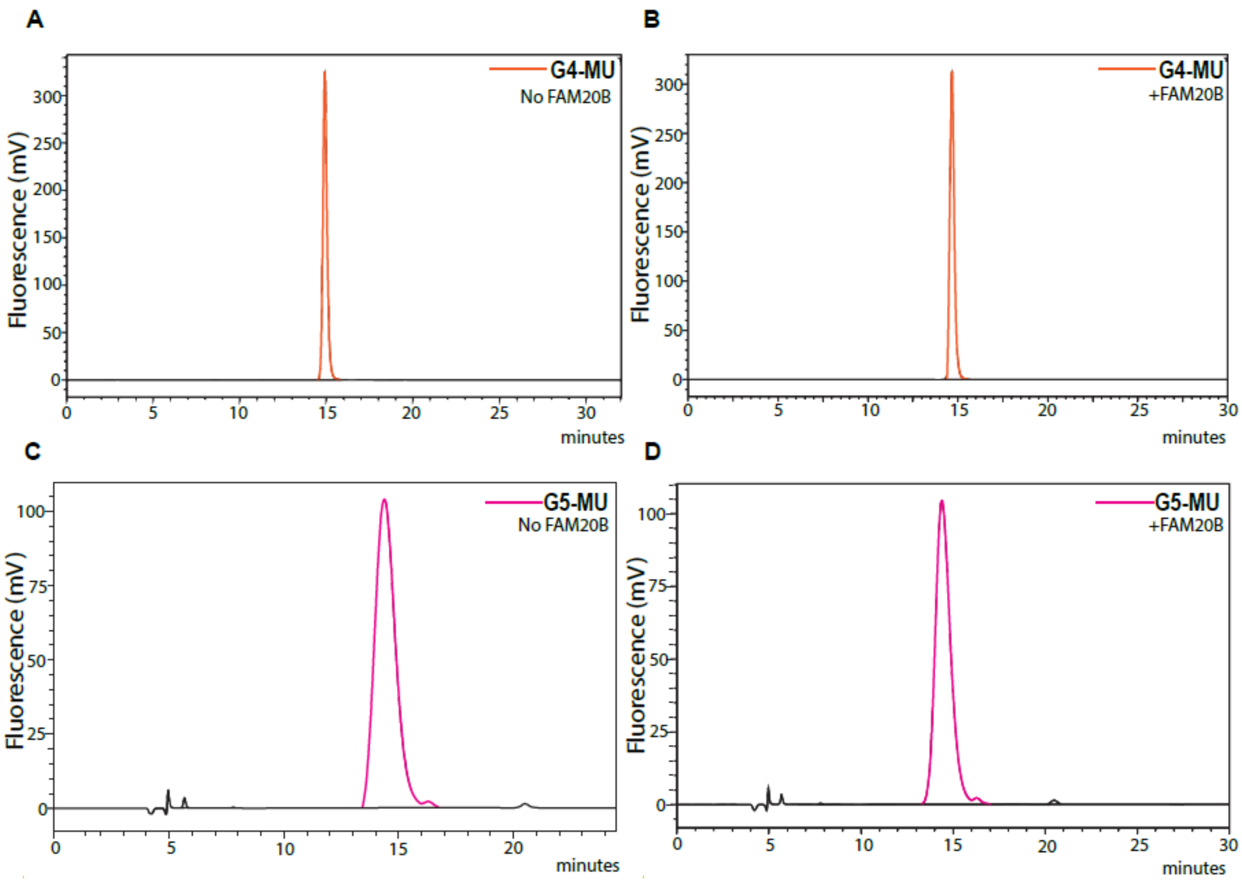
A
G4-MU
No FAM20B
Fluorescence (mV)
minutes
B
G4-MU
+FAM20B
Fluorescence (mV)
minutes
C
G5-MU
No FAM20B
Fluorescence (mV)
minutes
D
G5-MU
+FAM20B
Fluorescence (mV)
minutes

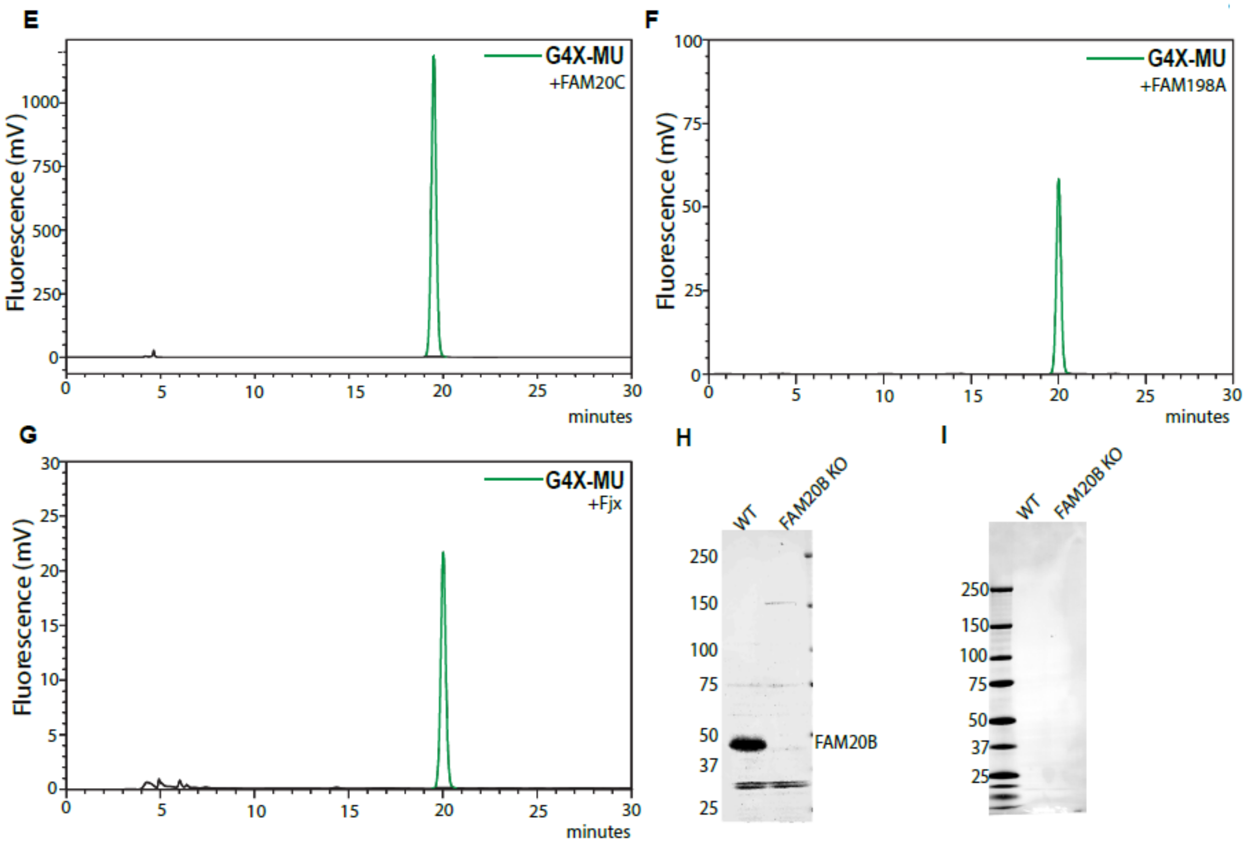


**Fig S1. FAM20B specifically phosphorylates the xylose of primer disaccharide.** (**A, B**) Chromatogram of G4-MU matriglycan (**A**) alone and (**B**) with FAM20B. 5-hour timepoint is shown. (**C, D**) Chromatogram showing G5-MU matriglycan (**C**) alone and (**D**) with FAM20B. (**E**) Chromatogram of primer disaccharide G4X-MU with FAM20C, 17-hour timepoint is shown. (**F**) Chromatogram of primer disaccharide G4X-MU with FAM198A, 16-hour timepoint is shown. (**G**) Chromatogram of primer disaccharide G4X-MU with Fjx-1, 16-hour timepoint is shown. (**H, I**) WT and FAM20B KO HAP cell lysates probed with (**H**) anti-Fam20B antibody and (**I**) anti-mouse IgG secondary antibody alone.

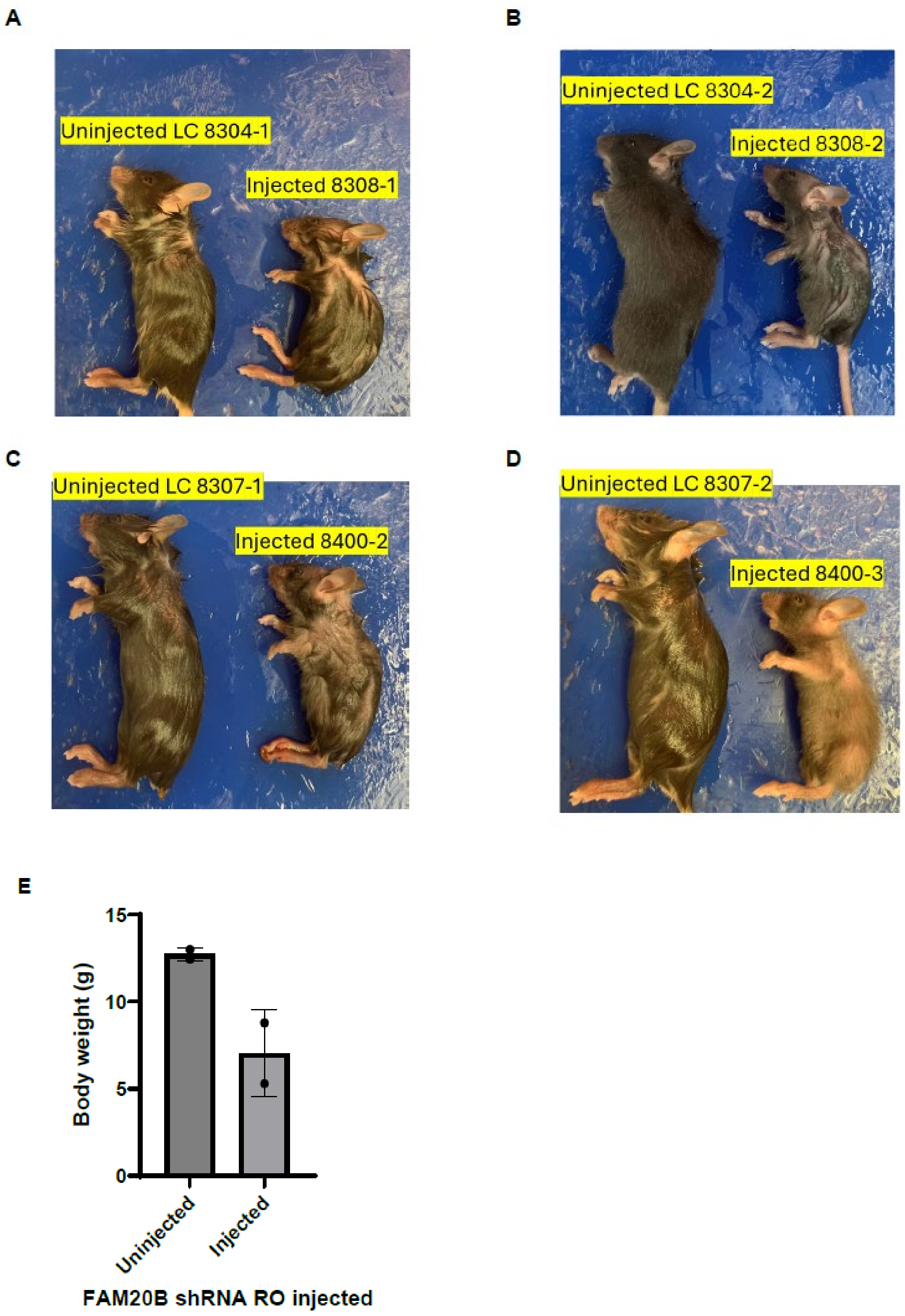
A
Uninjected LC 8304-1
Injected 8308-1
B
Uninjected LC 8304-2
Injected 8308-2
C
Uninjected LC 8307-1
Injected 8400-2
D
Uninjected LC 8307-2
Injected 8400-3
E
Body weight (g)
15
10
5
0
Uninjected
Injected
FAM20B shRNA RO injected

**Fig S2. FAM20B knockdown causes reduced body weight, stunted growth and early lethality in mice.** (**A**-**D**) Injected mice and their uninjected littermates are shown. (**E**) Body weights of injected versus uninjected mice.

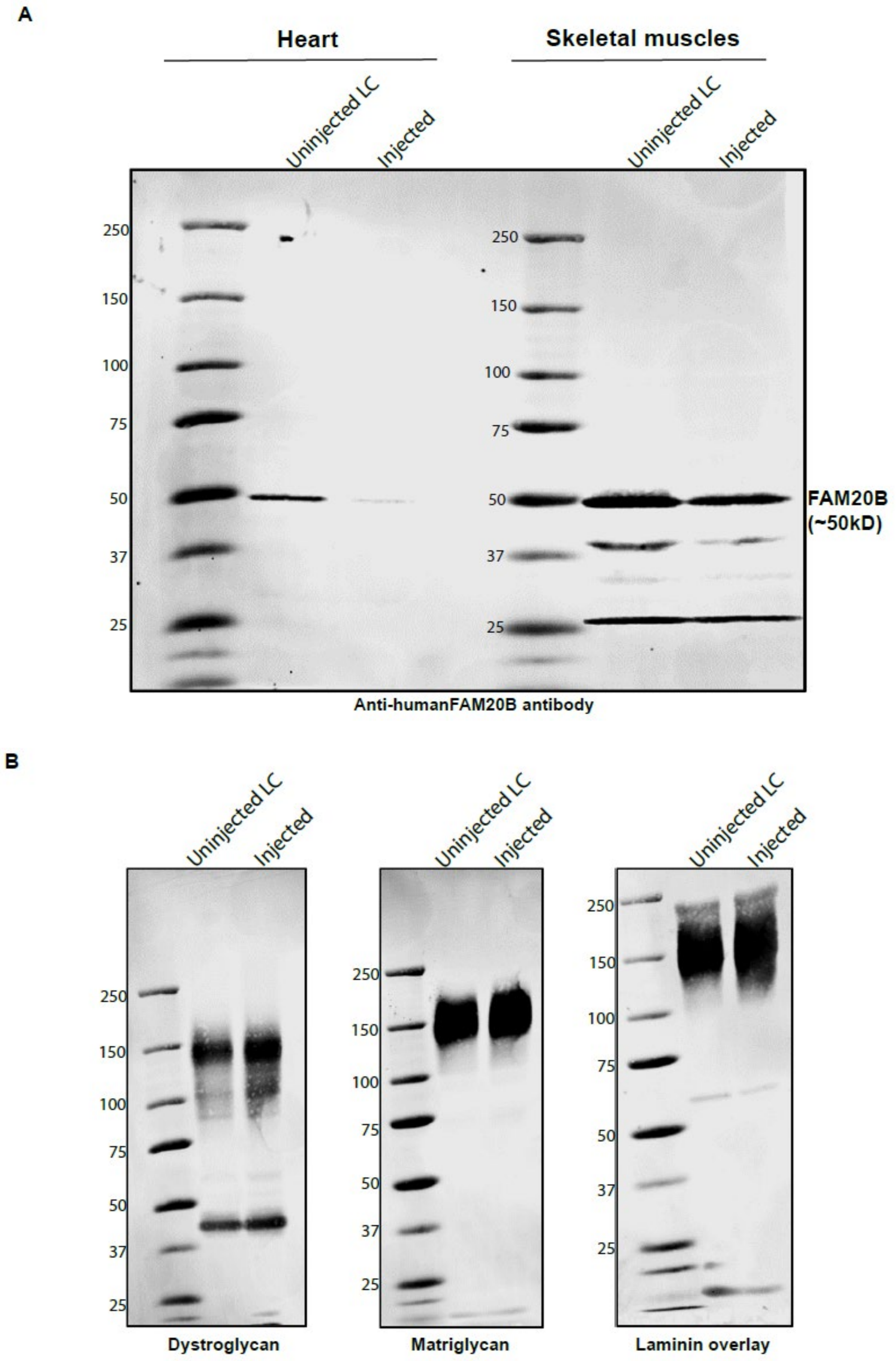


**Fig S3. FAM20B shRNA injections reduce levels of FAM20B protein in the heart of mice.**
(**A**) Cell lysates from heart and skeletal muscles of injected and uninjected littermate control

(LC) mice were probed with anti-human FAM20B antibody. Molecular weights are indicated on the left of each blot. (**B**) Immunoblot analysis of skeletal muscle from uninjected littermate control (LC) and injected mice. Glycoproteins were enriched using wheat-germ agglutinin (WGA)-agarose with 10 mM EDTA. Immunoblotting was performed to detect matriglycan (IIH6), core α-DG, β-DG (AF6868) (the broad 100-200 kDa band represents α-DG and the discrete ~ 42 kDa band represents β-DG), and laminin overlay. Molecular weight standards in kilodaltons (kDa) are indicated on the left of each blot.

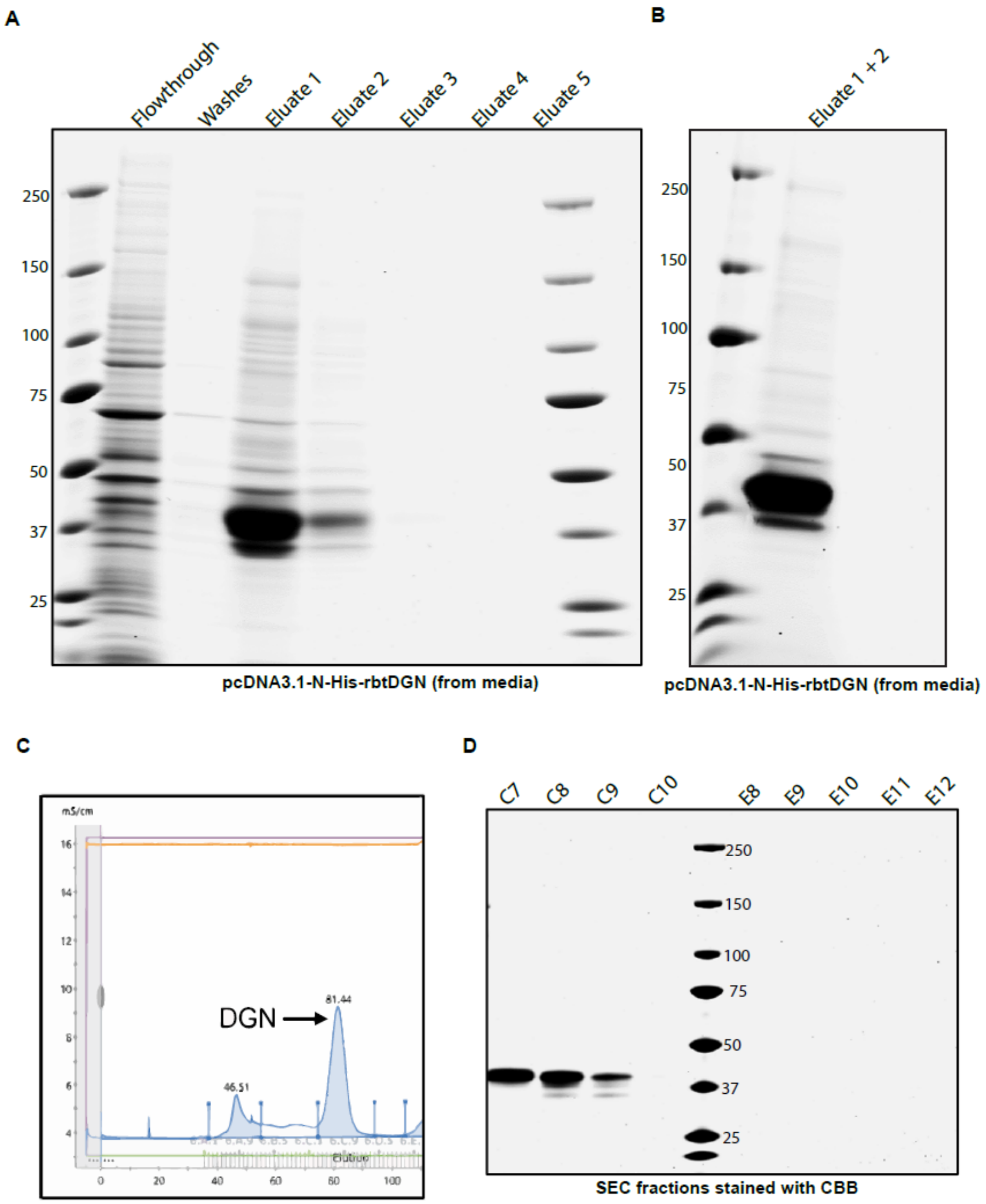
A
Flowthrough
Washes
Eluate 1
Eluate 2
Eluate 3
Eluate 4
Eluate 5
250
150
100
75
50
37
25
pcDNA3.1-N-His-rbtDGN (from media)
B
Eluate 1 + 2
250
150
100
75
50
37
25
pcDNA3.1-N-His-rbtDGN (from media)
C
mS/cm
DGN
81.44
46.51
Elution
D
C7
C8
C9
C10
E8
E9
E10
E11
E12
250
150
100
75
50
37
25
SEC fractions stained with CBB

E

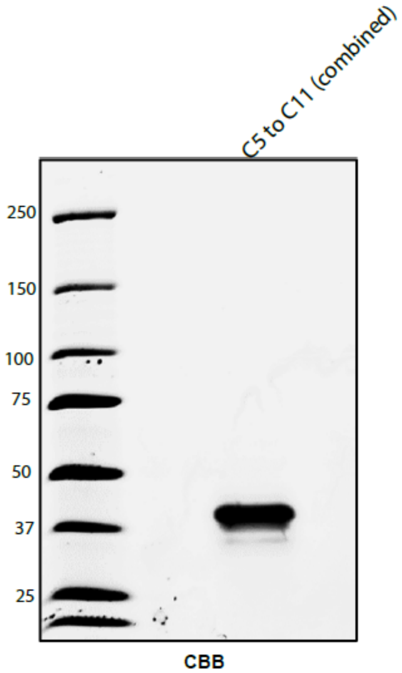


**Fig S4. Purification of recombinant and tagged N-terminal domain of dystroglycan (α-DGN) from mammalian cells.** (**A**) 6X-His tagged α-DGN was secreted in the media and purified using TALON metal affinity chromatography. SDS-PAGE stained with Coomassie Brilliant Blue (CBB) depicting the various purification steps is shown. α-DGN is approximately 37 kDa. Molecular weights are indicated on the left. (**B**) Relevant eluates containing α-DGN were combined, concentrated and buffer exchanged with 1X PBS pH 7.4. The combined eluates were run on an SDS-PAGE gel and stained with CBB are shown. Molecular weight standards in kilodaltons (kDa) are indicated on the left. (**C**) The combined eluate from TALON chromatography was subjected to size exclusion chromatography. Proteins were detected by UV absorbance at 280 nm. The large absorbance peak depicts α-DGN. (**D**) The relevant fractions containing α-DGN (~37 kDa) were collected, ran on an SDS-PAGE gel and stained with CBB. Molecular weights are indicated on the left. (**E**) Fractions confirmed to have α-DGN were again collected, combined, concentrated, ran on SDS-PAGE gel and stained with CBB. Molecular weight standards in kilodaltons (kDa) are indicated on the left.

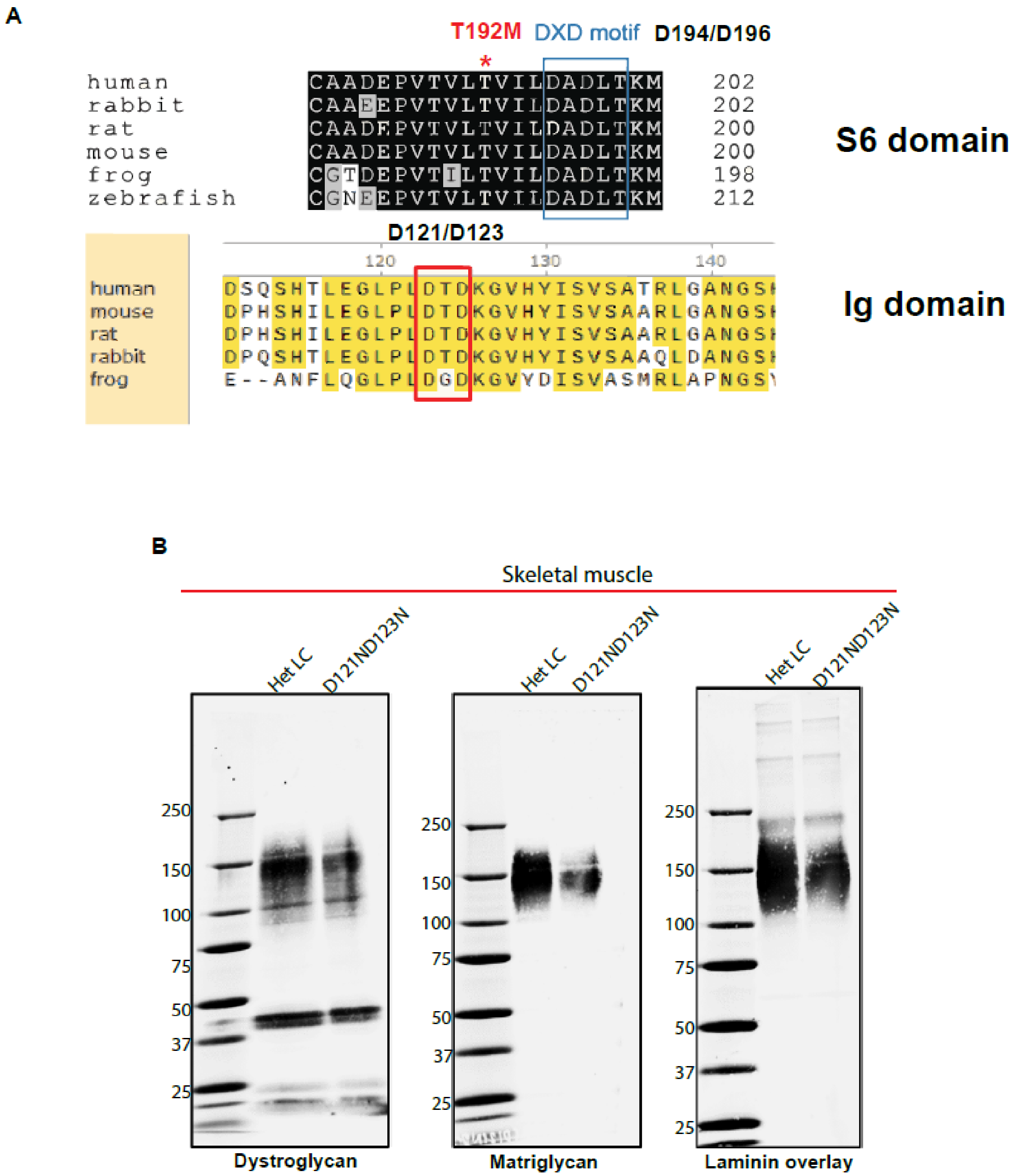
A
T192M
DXD motif
D194/D196
human CAADEPVTVLTVILDADLTKM 202
rabbit CAAEEPVTVLTVILDADLTKM 202
rat CAADEPVTVLTVILDADLTKM 200
mouse CAADEPVTVLTVILDADLTKM 200
frog CGTDEPVTILTVILDADLTKM 198
zebrafish CGNEEPVTVLTVILDADLTKM 212
S6 domain
D121/D123
120
130
140
human DSQSHTLEGLPLDTDKGVHYISVSATRLGANGS
mouse DPHSHILEGLPLDTDKGVHYISVSAARLGANGS
rat DPHSHILEGLPLDTDKGVHYISVSAARLGANGS
rabbit DPQSHTLEGLPLDTDKGVHYISVSAAQLDANGS
frog E--ANFLQGLPLDGDKGVYDISVASMRLAPNGS
Ig domain
B
Skeletal muscle
Het LC
D121ND123N
250
150
100
75
50
37
25
Dystroglycan
Matriglycan
Laminin overlay

C

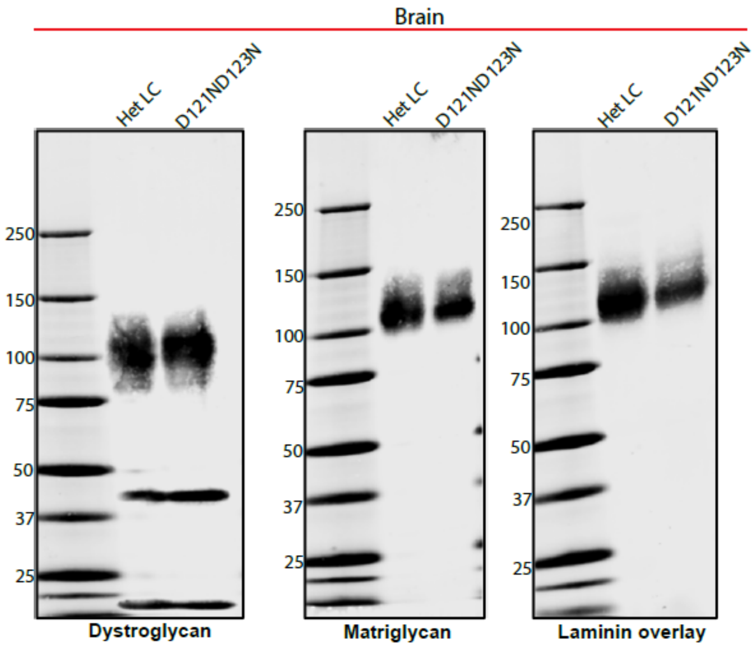
Brain
Het LC
D121ND123N
Het LC
D121ND123N
Het LC
D121ND123N
250
150
100
75
50
37
25
250
150
100
75
50
37
25
250
150
100
75
50
37
25
Dystroglycan
Matriglycan
Laminin overlay

D

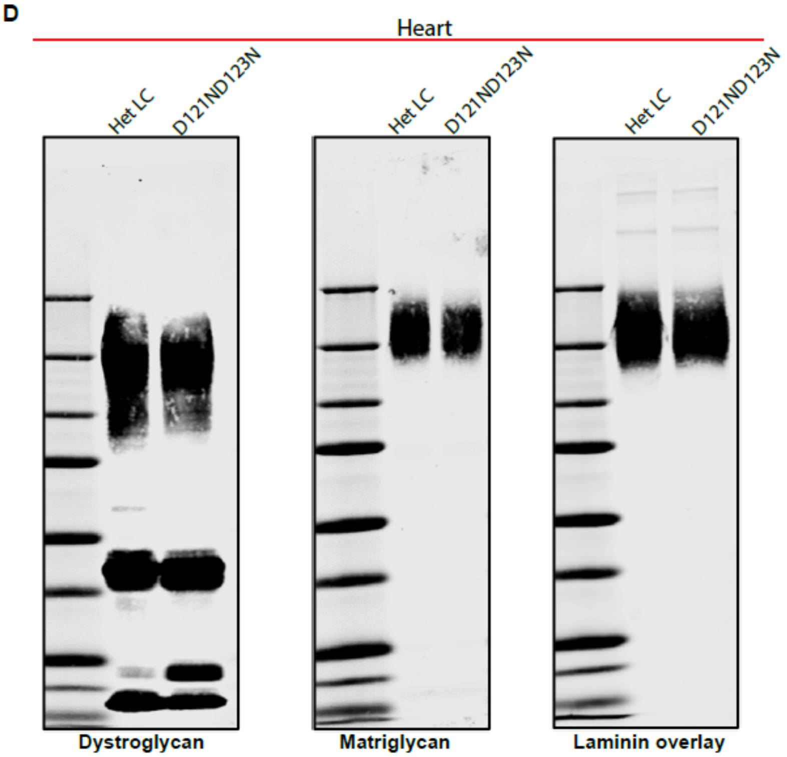
Heart
Het LC
D121ND123N
Het LC
D121ND123N
Het LC
D121ND123N
Dystroglycan
Matriglycan
Laminin overlay

E

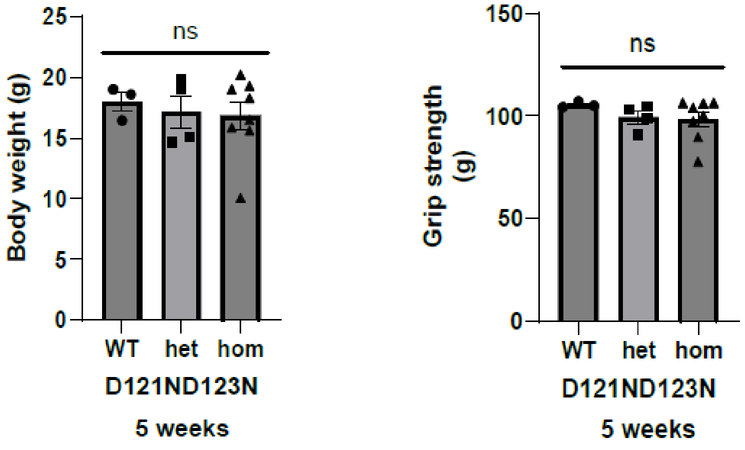


**Fig S5. D121N/D123N mutation in α-DGN does not change molecular weights of matriglycan, DG and maintains normal physiology in mice.** (**A**) Sequence comparison of the N-terminal domain of dystroglycan (DGN) from various species shows it has two DXD motifs, one in S6 domain and another in Ig-like domain**.** (**B-D**) Immunoblot analysis of (**B**) skeletal muscle, (**C**) brain and (**D**) heart from heterozygous littermate control (Het LC) and mutant mice. Glycoproteins were enriched using wheat-germ agglutinin (WGA)-agarose with 10 mM EDTA. Immunoblotting was performed to detect matriglycan (IIH6), core α-DG, β-DG (AF6868) (the broad 100-200 kDa band represents α-DG and the discrete ~ 42 kDa band represents β-DG), and laminin overlay. Molecular weight standards in kilodaltons (kDa) are indicated on the left of each blot. (**E**) Body weight and grip strength of 5-week-old wild-type (WT) littermate control and mutant heterozygous (het) and homozygous (hom) mutant mice. Statistical significance was determined using Ordinary one-way ANOVA with Tukey's post hoc test, ns = not significant,

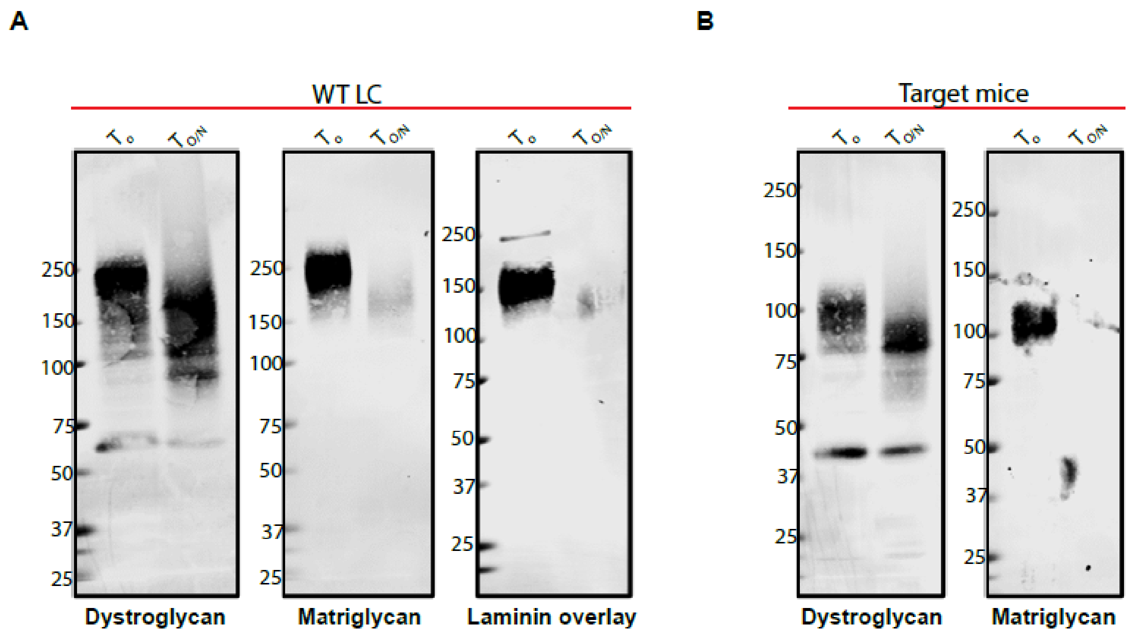


**Fig S6. The shorter band detected in D194N/D196N mutants is matriglycan. (A, B)** Immunoblot analysis of total skeletal muscle from (**A**) Wild-type (WT) littermate control (LC) mice (left) and (**B**) target mutant mice (right) after digestion with enzymes β-glucuronidase (BGUS) and α-xylosidase (XyIS). WGA enriched glycoproteins were incubated overnight with BGUS and XyIS. Immunoblotting was performed to detect matriglycan (IIH6), core α-DG and β-DG (AF6868), and laminin overlay before ($T_o$) and after overnight digestion ($T_{O/N}$). Molecular weight standards in kilodaltons (kDa) are shown on the left.

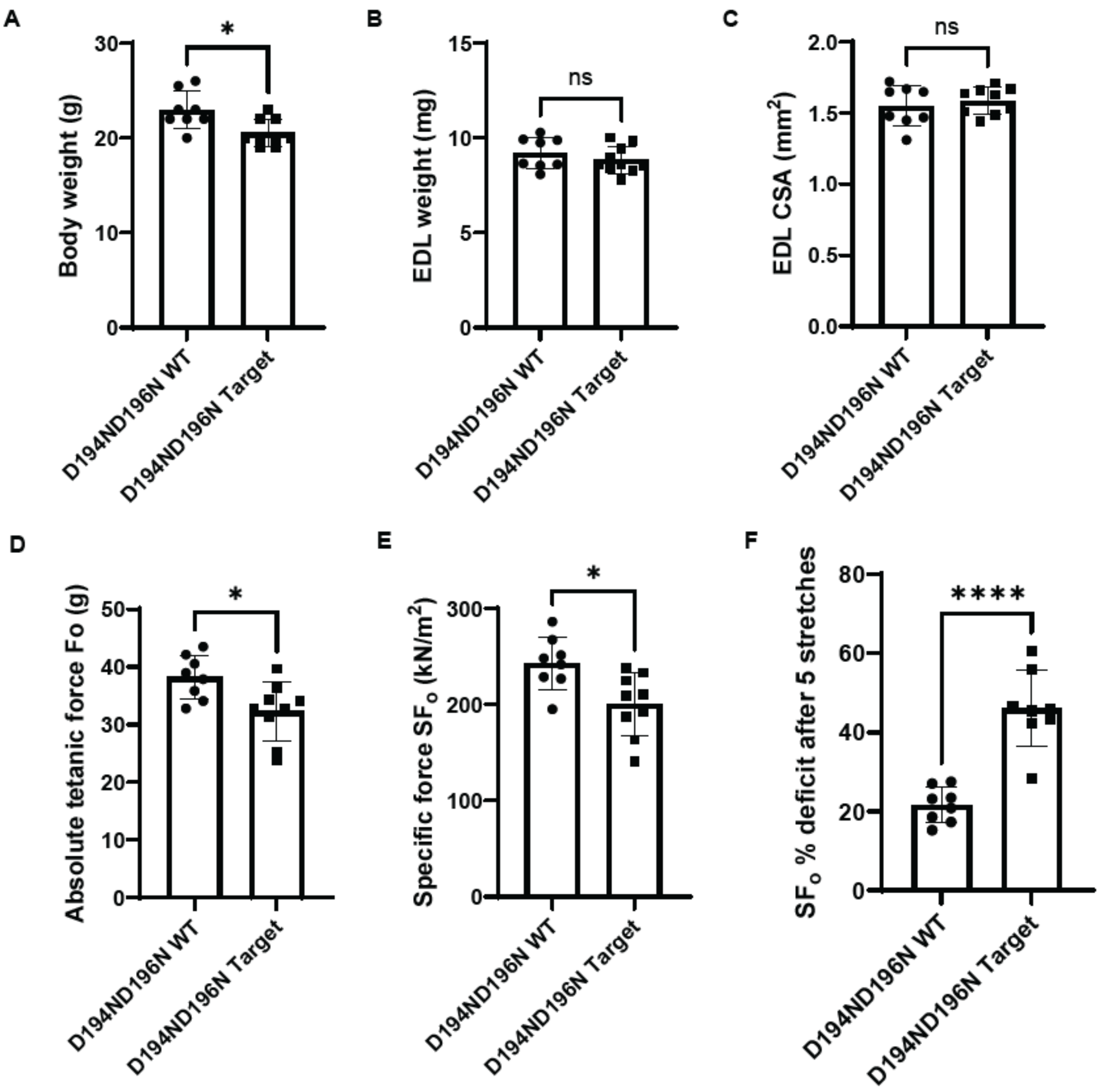
A
Body weight (g)
*
D194ND196N WT
D194ND196N Target
B
EDL weight (mg)
ns
C
EDL CSA (mm2)
ns
D
Absolute tetanic force Fo (g)
*
E
Specific force SFo (kN/m2)
*
F
SFo % deficit after 5 stretches
****

G

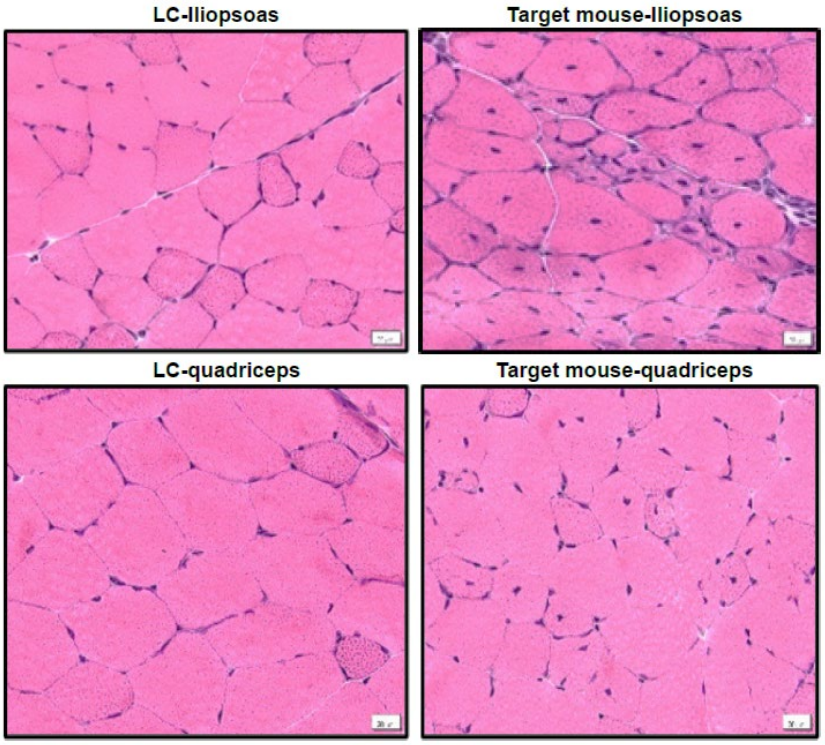


**Fig S7. The shorter matriglycan in D194N/D196N mutants is unable to maintain muscle function. (A-G)** Force production analysis of EDL muscle from 12-18 weeks old D194N/D196N female mice. (**A**) Body weights, (**B**) EDL weights, (**C**) cross sectional area (CSA) of EDL muscle, (**D**) absolute tetanic force, (**E**) specific force, (**F**) specific force deficit after 5 stretches and (**G**) histological analyses of iliopsoas and quadricep muscles from 6-week-old littermate control (LC) and target mice. Centrally nucleated muscle cells are visible in the target mice. Sections are stained with H&E. Scale bar 20µm.
